# Kindlin-2-Moesin interaction orchestrates sprouting angiogenesis via modulating endothelial membrane mechanics and VEGF signaling

**DOI:** 10.64898/2026.02.24.707842

**Authors:** Lu Wang, Yuxin Fu, Zeyang Yu, Yi Lei, Tianjing Yang, Jiayu Liu, Nina Ma, Yuming Liu, Kunfu Ouyang, Kai Zhang, Junhao Hu, Xi Fang, Ying Shen, Jing Zhou, Xiaohong Wang

**Author notes:** Corresponding author. (X.W) or (J.Z).

## Abstract

Membrane mechanics play a crucial role in cellular signaling and fate determination, yet their impact on angiogenesis remains poorly understood. Here, we identify Kindlin-2 as a key regulator of sprouting angiogenesis via regulating endothelial membrane tension through its interaction with Moesin, a crucial linker protein between the cell membrane and the actin cortex. Mechanistically, Kindlin-2 binds to the N62 residue of Moesin, limiting its overactivation and maintaining proper membrane tension to facilitate VEGFR2 endocytosis and downstream signaling. Using both developmental and pathological models, we demonstrate that the interaction of Kindlin-2 and Moesin is enhanced in high angiogenic conditions, and endothelial Kindlin-2 deletion reduces angiogenesis. Furthermore, mutation of Moesin at N62 phenocopies the effects of Kindlin-2 loss. Together, these findings uncover a previously unrecognized mechanism linking membrane tension regulation to angiogenesis and provide new insights into targeting the Kindlin-2–Moesin axis for therapeutic intervention in neovascular diseases.

## Introduction

The blood vascular network is a highly branched system that supplies all organs in vertebrates, playing a critical role in development, physiological homeostasis, and wound healing. Its dysfunction is implicated in various diseases, including solid tumors, neovascular ocular disorders, obesity, and others (*1*). A major focus of angiogenesis research is the effect of vascular endothelial growth factor (VEGF) on blood vessels. VEGF regulates key aspects of vascularization, such as endothelial cell (EC) sprouting, migration, proliferation, and survival (*2*). Furthermore, vascular cells exhibit mechanosensitivity, and external mechanical forces are increasingly recognized as essential regulators of vascular functions (*3*).

Organ development involves processes such as growth, movement, and tissue reorganization, which generate mechanical forces that cells can perceive. In the vascular system, mechanical cues such as fluid shear stress (*4, 5*), extracellular matrix (ECM) stiffness (*6, 7*), and cyclic stretch (*8, 9*) regulate vascular development, homeostasis and pathologies. While most studies have focused on how extracellular mechanical factors influence EC function, the interplay between cell-intrinsic mechanics and molecular signaling in driving sprouting angiogenesis remains largely unexplored.

Kindlins are cytoskeleton-associated proteins best known for their role in binding to the cytoplasmic tails of β-integrin, where they regulate integrin activation, cell adhesion, and signaling. Among the three Kindlin family members (Kindlin-1, -2, and -3), Kindlin-2 is ubiquitously expressed, whereas Kindlin-1 and -3 exhibit more restricted expression patterns, primarily in epithelial and hematopoietic cells, respectively (*10*). Mice with a global knockout of *Kindlin-2* exhibit peri-implantation lethality by embryonic day 7.5 (E7.5) due to defects in endodermal and epiblast attachment, underscoring its critical role in early development (*11, 12*). While much of the research on Kindlin-2 has centered on its contributions to tumor formation, progression, and metastasis (*13*), growing evidence suggests that Kindlin-2 also regulates organogenesis and homeostasis through both integrin-dependent and integrin-independent mechanisms (*14–16*). Additionally, Kindlin-2 has emerged as a mechano-responsive protein capable of responding to mechanical cues such as tension, ECM stiffness, and shear stress (*13, 17, 18*). In ECs, Kindlin-2 is highly expressed and plays a key role in maintaining vascular stability in response to shear stress (*18*). It is also essential for vascular development, as *Tie2*-Cre-mediated deletion of *Kindlin-2* leads to embryonic lethality as early as E10.5 (*19*). However, how Kindlin-2 regulates sprouting angiogenesis and the underlying mechanisms remain elusive.

The cell surface is a mechanobiological unit which compass the plasma membrane, a lipid bilayer embedded with proteins, and the underlying actomyosin-rich cortical cytoskeleton, also known as the cell cortex (*20*). Plasma membrane tension, defined as the in-plane resistive force opposing surface expansion, plays a critical role in transmitting and integrating cellular and tissue-level signals. This tension is largely regulated by membrane-cortex attachment (MCA) (*21–23*). MCA proteins, including members of the ezrin-radixin-moesin (ERM) family, serve as physical linkers between the plasma membrane and the actin cortex, which provide resistance to the expansion of the membrane thereby regulating membrane tension. Among the three ERM proteins, Moesin is highly enriched in ECs (*24*) and has been implicated in various EC functions, including motility, angiogenesis, and vascular permeability (*25, 26*). However, the mechanisms by which Moesin regulates EC behavior and how its activity is controlled remain largely unclear.

Here, we show that Kindlin-2 is essential for sprouting angiogenesis. Endothelial-specific loss of Kindlin-2 leads to severe vascular developmental defects, including impaired sprouting and proliferation. Mechanistically, we identify Kindlin-2 as a previously unrecognized regulator of Moesin, interacting with Moesin to prevent its overactivation. This interaction is crucial for maintaining appropriate membrane tension, facilitating vascular endothelial growth factor receptor type 2 (VEGFR2) endocytosis, and ensuring effective VEGF signal transduction. Together, our findings provide mechanistic insight into how intrinsic membrane mechanics are coupled with molecular signaling pathways to regulate sprouting angiogenesis.

## Results

### Endothelial Kindlin-2 is required for sprouting angiogenesis

To investigate the role of Kindlin-2 in blood vessels, we first examined its expression in the mouse retina, a well-established model for studying sprouting angiogenesis. Immunofluorescence staining showed that at postnatal day 6 (P6), Kindlin-2 was highly expressed in retinal blood vessels, with elevated expression at the vascular front compared to the vascular plexus **(Fig. S1A)**. Analysis in adulthood also revealed Kindlin-2 is highly expressed in the retinal vasculature, while the expression seems lower compared with developing retina (**Fig. S1B**). Additionally, publicly available single-cell RNA-seq data (GSE175895) also showed that *Fermt2* (encoding Kindlin-2) is enriched in ECs in P10 retina (**Fig. S1C**).

To explore the functional role of Kindlin-2 in growing blood vessels, we generated endothelial-specific *Kindlin-2* knockout mice (*Kindlin-2^ΔEC(Tie2)^*) by crossing *Kindlin-2^fl/fl^* mice (*27*) with *Tie2-Cre* mice. Consistent with previous reports, loss of Kindlin-2 in ECs resulted in embryonic lethality around embryonic day 10.5 (E10.5) (**Fig. S1D and E**). To circumvent this early lethality, we generated inducible EC-specific *Kindlin-2* knockout mice (*Kindlin-2^iΔEC(Pdgfb)^*) by crossing *Kindlin-2*^fl/fl^ mice with *Pdgfb-CreERT2* mice (*28*). Tamoxifen was administered from E8.5 to E10.5, and embryos were analyzed at E12.5 (**Fig. 1A**). *Kindlin-2^iΔEC(Pdgfb)^*embryos exhibited severe hemorrhaging, and analysis of brain vasculature revealed a marked reduction in vascularized area compared to controls (**Fig. 1B and C**).

**Figure 1.**
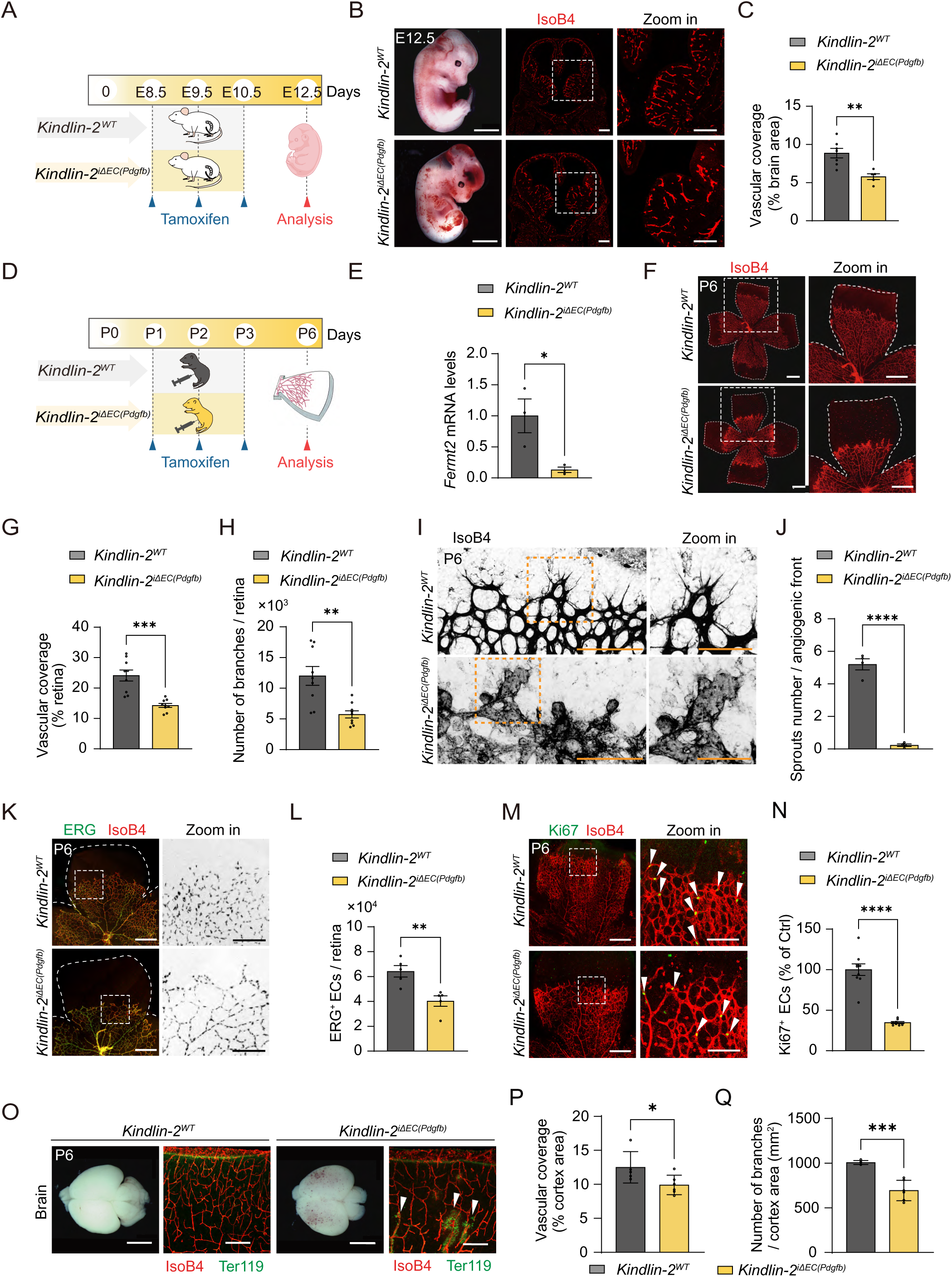
Endothelial Kindlin-2 is required for sprouting angiogenesis. (A) Schematic illustration of tamoxifen administration in pregnant mice. (B) Left: representative images of E12.5 *Kindlin-2^WT^* and *Kindlin-2^iΔEC(Pdgfb)^* embryos. Middle and right: representative confocal images of E12.5 embryonic brain sections stained with IsoB4 (red). (C) Quantification of vascular coverage in the brain shown in (D) (n = 7/5 embryos). (D) Schematic of tamoxifen administration schedule in *Kindlin-2^WT^*and *Kindlin-2^iΔEC(Pdgfb)^* mice. Tamoxifen was administered from P1 to P3, and retinas were analyzed at P6. (E) Quantification of *Fermt2* mRNA (encode Kindlin-2) levels in brain endothelial cells (ECs) isolated from P6 *Kindlin-2^iΔEC(Pdgfb)^*pups (n = 3/3 pups). (F) Representative confocal images of flat-mounted retinas from P6 *Kindlin-2^WT^* and *Kindlin-2^iΔEC(Pdgfb)^* pups stained with IsoB4 (red). (G-H) Quantification of (H), showing reduced vascular area and branch points in *Kindlin-2^iΔEC(Pdgfb)^* pups (n = 9/9 pups). (I) Representative images of the angiogenic front in P6 *Kindlin-2^WT^* and *Kindlin-2^iΔEC(Pdgfb)^* retinas stained with IsoB4. Higher-magnification images show blunted morphology of tip cells. (J) Quantification of (K), showing a reduced number of sprouts in *Kindlin-2^iΔEC(Pdgfb)^* pups (n = 4/4 pups). (K) Representative confocal images of ERG (EC nuclei, green) and IsoB4 (red) in flat-mounted retinas from P6 *Kindlin-2^WT^* and *Kindlin-2^iΔEC(Pdgfb)^*pups. (L) Quantification of ERG^+^ nuclei in (M) (n = 5/5 pups). (M) Representative confocal images of Ki67 (green) and IsoB4 (red) in flat-mounted retinas from P6 *Kindlin-2^WT^* and *Kindlin-2^iΔEC(Pdgfb)^*pups. (N) Quantification of Ki67^+^ cells in the vascular area of *Kindlin-2^WT^*and *Kindlin-2^iΔEC(Pdgfb)^* pups (n = 9/8 pups). (O) Representative images of brains from P6 *Kindlin-2^WT^* and *Kindlin-2^iΔEC(Pdgfb)^*pups and immunofluorescence staining of Ter119 (erythrocytes, green), IsoB4 (red) in brain sections. White arrows indicate hemorrhage area. (P-Q) Quantification of (Q), showing reduced vascular coverage and branch points in *Kindlin-2^iΔEC(Pdgfb)^* brains (n = 5/6 pups). Data are presented as mean ± SEM. \**P* < 0.05; ** *P* < 0.01; *** *P* < 0.001; **** *P* < 0.0001 by two-tailed Student’s t-test. Scale bars: 1 mm in brightfield panels of (B) and (O); 500 μm in (B), (F), and low magnification of (K), and (M); 200 μm in low magnification of (I), and high magnification of (K), and (M); 100 μm in (O) and high magnification of (I).

Next, we assessed the role of Kindlin-2 in sprouting angiogenesis in the developing mouse retina. Cre recombination was induced in newborn pups from P1 to P3, as previously described, (*29*), and the pups was analyzed at P6 (**Fig. 1D**). The knockout efficiency was confirmed in isolated brain ECs (**Fig. 1E**). No significant differences in body weight or the weight of major organs including brain, heart, lung, liver, and kidney were observed at P6 (**Fig. S1F-K**). Whole-mount retinal analysis revealed a significant reduction in vascular coverage and branch points in *Kindlin-2^iΔEC(Pdgfb)^* pups compared to littermate controls (**Fig. 1F-H**). Strikingly, tip ECs in the vascular front region of *Kindlin-2^iΔEC(Pdgfb)^* mice exhibited a blunted, aneurysm-like structure, with fewer and dysmorphic filopodia (**Fig. 1I and J**). In addition, the total number of ECs, as determined by counting ERG^+^ nuclei per retina, as well as the number of proliferating ECs (Ki67^+^ IsoB4^+^ cells), were also reduced in *Kindlin-2 ^iΔEC(Pdgfb)^* retinas (**Fig. 1K-N**). Similarly, the brain of *Kindlin-*2*^iΔEC(Pdgfb)^*group exhibited hemorrhages, with reduced vascular coverage and branch points (**Fig. 1O-Q**).

The formation of the deep retinal vasculature begins at P7 and completes by P14 (*30*). To determine whether Kindlin-2 also regulates deep vascular network formation, Cre recombination was induced between P4-P6, and mice were analyzed at P9 (**Fig. S2A**). No significant differences in body weight or major organ weights were observed (**Fig. S2B-G**). Similar to the superficial retinal vascular results, loss of Kindlin-2 markedly impaired the formation of the deep vascular plexus (**Fig. S2H-L**). Consistent with this, vascular coverage and number of branch points were also significantly reduced in the brain cortex of *Kindlin-2^iΔEC(Pdgfb)^* pups at P9 (**Fig. S2M-O**).

To assess whether the vascular defects in the retina lead to visual impairment, Cre recombination was induced at P14, and electroretinography (ERG) was performed in P21 *Kindlin-*2*^iΔEC(Pdgfb)^*and *Kindlin-2^WT^* mice (**Fig. S2P**). Both A-wave and B-wave amplitudes were significantly decreased in *Kindlin-*2*^iΔEC(Pdgfb)^* mice compared to *Kindlin-2^WT^* littermates, indicating substantial visual impairment in *Kindlin-2*-deficient mice (**Fig. S2Q-S**).

To further confirm the phenotype observed in *Pdgfb-CreERT2*-mediated *Kindlin-2* knockout, we generated another inducible EC-specific *Kindlin-2* knockout model by crossing *Kindlin-2^fl/fl^* mice with *Cdh5-(PAC)-CreERT2* mice (termed *Kindlin-2^iΔEC(Cdh5)^*). Tamoxifen was administered from P1 to P3, and retinal angiogenesis was analyzed at P6 (**Fig. S3A**). Reduced vascular coverage, reduced branching, and fewer number of sprouts were observed in *Kindlin-2^iΔEC(Cdh5)^* pups, similar to the *Pdgfb-CreERT2*-mediated *Kindlin-2* EC-specific knockout (**Fig. S3B-F**).

By using an ex vivo aortic ring assay we further showed that the vascularization and sprouting were significantly reduced in *Kindlin-2^iΔEC(Pdgfb)^*aortic ring (**Fig. S3G and H**). To further investigate the role of endothelial Kindlin-2 in filopodia formation, we employed an in vitro model which is the wound scratch-induced filopodia formation assay (*31*). Human umbilical vein endothelial cells (HUVECs) were transfected with *KINDLIN-2* siRNA, and the knockdown efficiency was confirmed by western blotting (**Fig. S3I and J**). We then compared filopodia formation in siCtrl-ECs and si*KINDLIN-2*-ECs after scratch wounding and VEGF stimulation. The results showed that si*KINDLIN-2*-ECs exhibited fewer phalloidin^+^ filopodia at the wound edge compared to siCtrl-ECs, in both unstimulated and VEGF-stimulated conditions. Notably, VEGF treatment increased the number of phalloidin^+^ filopodia at the leading edge in siCtrl-ECs, whereas *KINDLIN-2* knockdown cells failed to do so (**Fig. S3K and L**). Since we observed reduced EC proliferation in the *Kindlin-*2*^iΔEC(Pdgfb)^*retinas **(Fig. 1O and P)**, we further performed a BrdU incorporation assay to assess proliferation in siCtrl- and si*KINDLIN-2*-transfected HUVECs following VEGF stimulation. Consistent with the filopodia results, *KINDLIN-2* knockdown reduced the number of BrdU^+^ cells under both basal and VEGF-stimulated conditions, and these cells failed to increase proliferation in response to VEGF (**Fig. S3M and N**). Together, these results suggest that endothelial Kindlin-2 is essential for sprouting angiogenesis and is required for EC responsiveness to VEGF.

### Loss of Kindlin-2 and Integrin β1 in ECs showed distinct phenotype

We next sought to investigate the underlying mechanism by which Kindlin-2 regulates sprouting angiogenesis. Since Kindlin-2 is primarily recognized as a β-Integrin-binding protein involved in Integrin activation and signaling, we first wondered whether Kindlin-2 promotes angiogenesis through Integrin activation. Integrin β1-subunit is acknowledged as a prominent member of β-Integrin in postnatal retinal vasculature (*32*). Indeed, gene expression analysis in human retinal endothelial cells (HRECs), HUVECs, human cortical microvessels endothelial Cells (HCMECs), and purified P5 mouse brain ECs showed that of *ITGB1* (encoding Integrin β1) is substantially higher than other Integrin beta (β) subunits across these endothelial populations (**Fig. S3O**). Based on this, we generated inducible EC-specific Integrin β1 knockout mice (*Itgb1^iΔEC(Pdgfb)^*) by crossing *Itgb1^fl/fl^* mice with *Pdgfb-CreERT2* mice and induced Cre recombination following the same protocol used for *Kindlin-2^iΔEC(Pdgfb)^*mice (**Fig. 2A**). Consistent with a previous study (*32*), we found that although the radial outgrowth of the retina was reduced in *Itgb1^iΔEC(Pdgfb)^,* the tip ECs sprouted normally (**Fig. 2B-E**). This phenotype differs markedly from that observed in *Kindlin-2^iΔEC(Pdgfb)^* mice, where tip cells displayed a blunted, aneurysm-like structure with fewer and dysmorphic filopodia (**Fig. 2D and E**). Furthermore, *Itgb1^iΔEC(Pdgfb)^* mice exhibited decreased vascular coverage and arterial branch points but increased coverage and branching in the perivenous vascular plexus (*32*). In contrast, *Kindlin-2^iΔEC(Pdgfb)^*mice showed reduced arterial branch points and vascular coverage in both the arterial and venous regions (**Fig. 2F-J**). These phenotypic differences suggest that Kindlin-2 regulates sprouting angiogenesis through mechanisms distinct from integrin β1 activation.

**Figure 2.**
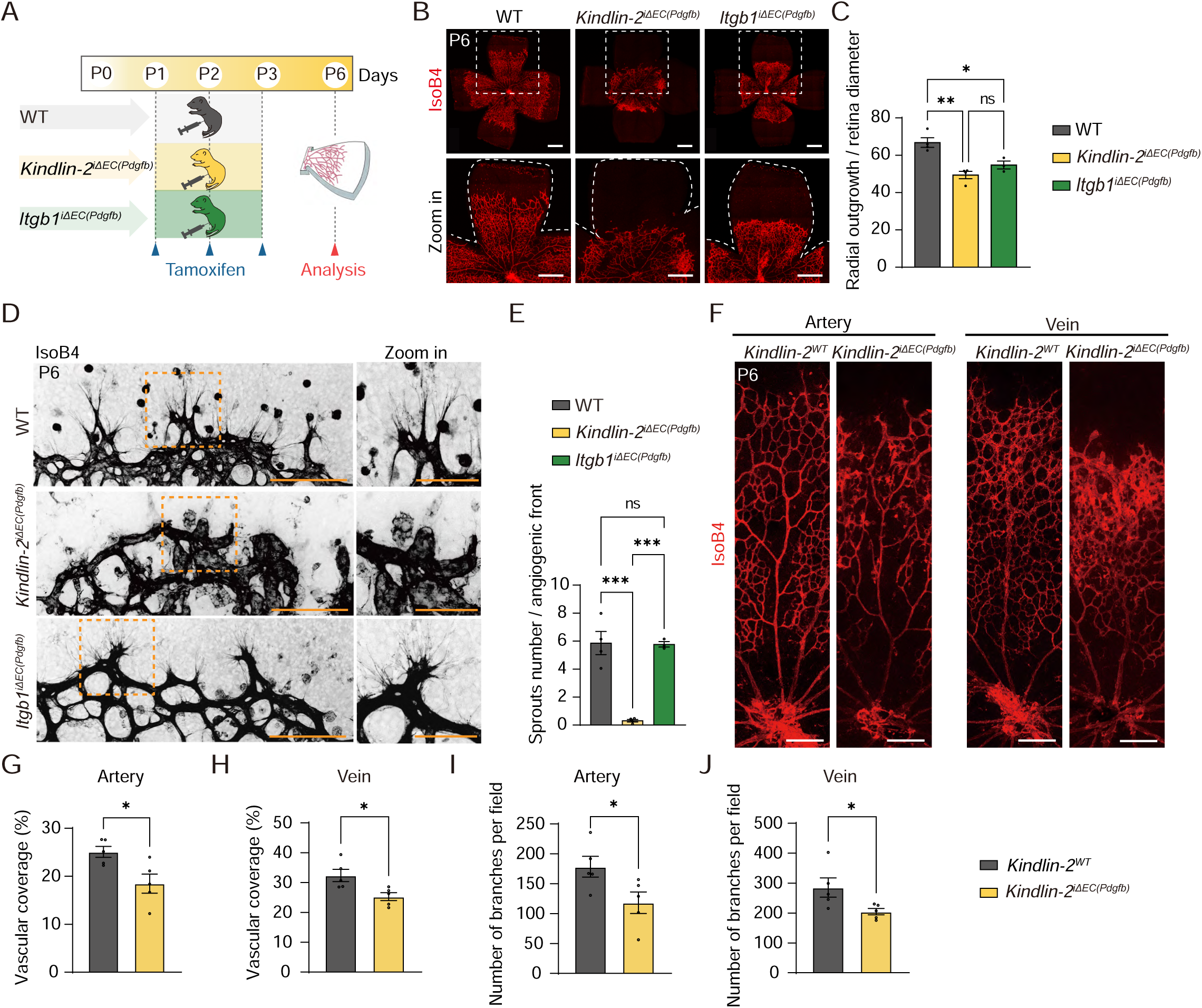
Loss of *Kindlin-2* or *Itgb1* in ECs results in distinct angiogenic phenotypes. (A) Schematic illustration of tamoxifen administration in WT, *Kindlin-2^iΔEC(Pdgfb)^*, and *Itgb1^iΔEC(Pdgfb)^* mice. Tamoxifen was administered from P1 to P3, and retinas were analyzed at P6. (B) Representative confocal images of flat-mounted P6 retinas from WT, *Kindlin-2^iΔEC(Pdgfb)^*, and *Itgb1^iΔEC(Pdgfb)^* pups stained with IsoB4 (red). (C) Quantification of radial vascular outgrowth in (B) (n = 4/4/3 pups). (D) Representative images showing the angiogenic front in P6 retinas of WT, *Kindlin-2^iΔEC(Pdgfb)^*, and Itgb1*^iΔEC(Pdgfb)^* mice. (E) Quantification of the number of sprouts in (D) (n = 4/3/3 pups). (F–J) Representative images and quantification of vascular coverage and branch points around arteries and veins in P6 retinas from *Kindlin-2^WT^* and *Kindlin-2^iΔEC(Pdgfb)^* pups (n = 5/5 pups). Data are presented as mean ± SEM. * *P* < 0.05; ** *P* < 0.01; *** *P* < 0.001 by two-tailed Student’s t-test or one-way ANOVA followed by Tukey’s multiple comparisons test. Scale bars: 500 μm in (B); 200 μm in (G) and low magnification of (E); 100 μm in high magnification of (E).

### Kindlin-2 associates with Moesin in ECs

Next, we aimed to identify potential binding partners of Kindlin-2 in ECs. For this, we expressed Ad-Flag-Kindlin-2 in cultured HUVECs and pulled down Kindlin-2 associated proteins using anti-Flag magnetic beads, followed by mass spectrometry analysis (**Fig. 3A**). Top five proteins ranked by the score were shown in the table **(Fig. 3B)**. Notably, Moesin was identified as a primary, previously unrecognized binding partner of Kindlin-2. Integrin β1 was also detected, though its binding efficiency was much lower compared with Moesin (**Fig. 3B**).

**Figure 3.**
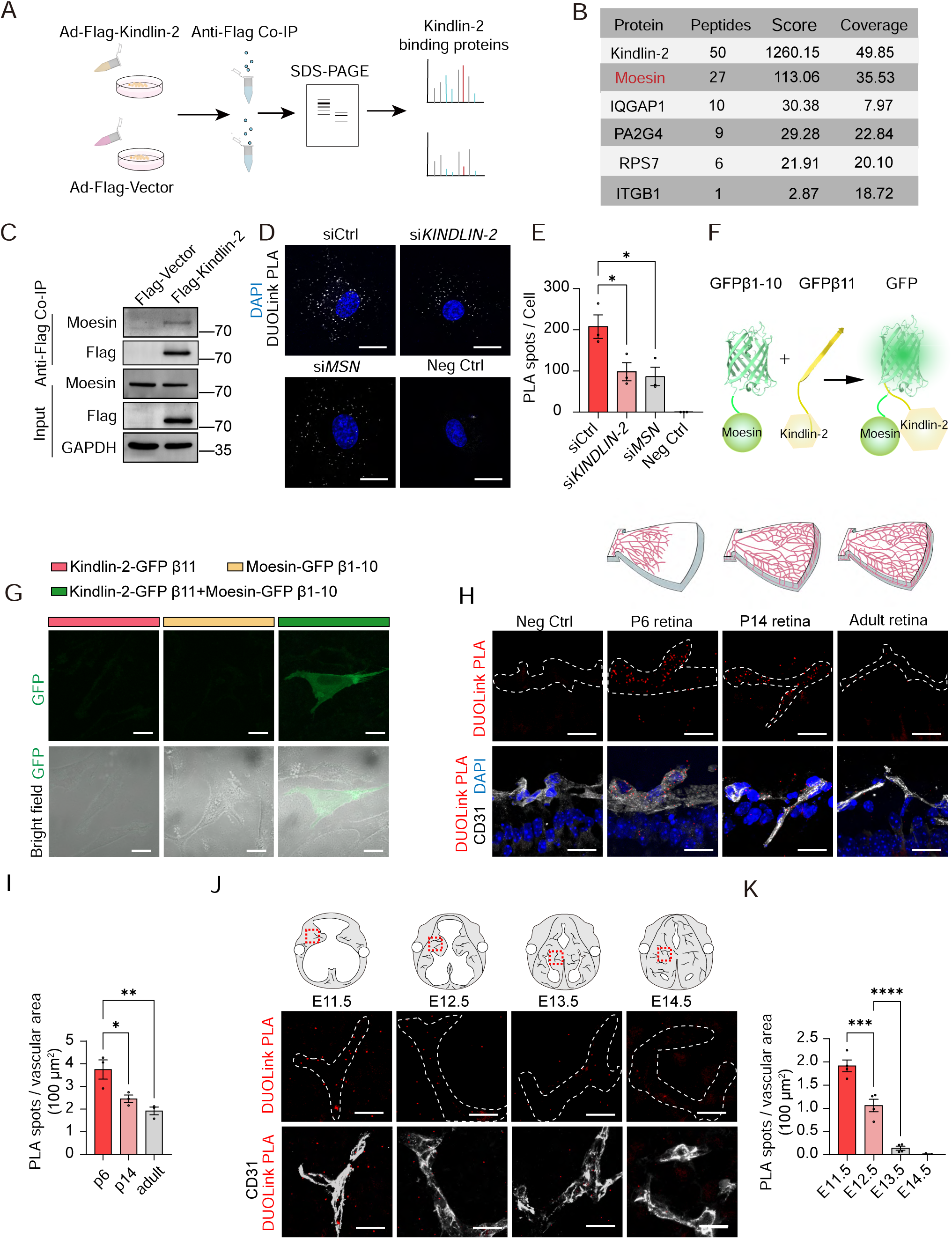
Kindlin-2 associates with Moesin in ECs. (A) Schematic illustration of the experimental design. Human umbilical vein endothelial cells (HUVECs) were infected with Ad-Flag-Kindlin-2 or Ad-Flag-Vector. Protein complexes were isolated from whole-cell lysates using anti-Flag M2 beads and analyzed by mass spectrometry. (B) Mass spectrometry results: the top five proteins ranked by the score and Integrin β1 were shown as binding proteins of Kindlin-2. (C) Co-IP analysis of Kindlin-2 and Moesin in HUVECs infected with Ad-Flag-Kindlin-2 or Ad-Flag-Vector. Moesin was pulled down with anti-Flag M2 beads and detected by western blot. Ad-Vector-infected cells served as control. (D) Representative images from proximity ligation assay (PLA) detecting Kindlin-2–Moesin interactions in HUVECs. Nuclei were stained with DAPI (blue), and PLA signals (white) indicate Kindlin-2–Moesin interactions. si*KINDLIN-2* (targeting Kindlin-2), si*MSN* (targeting Moesin), and omission of primary antibodies were used as negative controls. (E) Quantification of PLA signal spots per nucleus in (D) (n = 3 independent experiments). (F) Schematic of the split-GFP complementation assay. Kindlin-2 was fused to GFP β-strand 11 (residues 215–230), and Moesin was fused to GFP β-strands 1–10 (residues 1–214). Interaction between the two proteins reconstitutes GFP fluorescence. (G) Representative images of split-GFP fluorescence in EA.hy926 cells, indicating Kindlin-2–Moesin interaction in ECs. (H) Representative images of PLA for Kindlin-2 and Moesin in retina sections at P6, P14, and adulthood. Blood vessels were stained with CD31 (white), nuclei with DAPI (blue), and PLA signals are shown in red. White dotted line indicates the CD31^+^ area. (I) Quantification of PLA signal spots in CD31^+^ vessels from (H) (n = 3 independent experiments). (J) Representative images of PLA for Kindlin-2 and Moesin in embryonic brain sections at E11.5, E12.5, E13.5, and E14.5. Blood vessels were stained with CD31 (white) and PLA signals are shown in red. White dotted line indicates the CD31^+^ area. (K) Quantification of PLA signal spots in CD31^+^ vascular areas in (J) (n = 4 independent experiments). Data are presented as mean ± SEM. \**P* < 0.05; ** *P* < 0.01, *** *P* < 0.001, **** *P* < 0.0001 by two-tailed Student’s t-test or one-way ANOVA followed by Tukey’s multiple comparisons test. Scale bars: 5 μm in (D), (G), (H), and (J).

To validate the interaction between Kindlin-2 and Moesin, we immunoprecipitated Flag-tagged Kindlin-2 from HUVECs and detected co-precipitation of Moesin (**Fig. 3C**). Conversely, immunoprecipitation of Flag-tagged Moesin confirmed that Kindlin-2 co-precipitated with Moesin (**Fig. S4A**). Additionally, we utilized in situ proximity ligation assay (PLA), a technique that detects protein-protein interactions in situ (within <40 nm distance) at endogenous protein levels. The results further validated the interaction between Moesin and Kindlin-2 in ECs (**Fig. 3D and E**). To complement this, we employed a Split GFP assay (*33*). We linked GFPβ1-10 with Moesin and GFPβ11 with Kindlin-2, and GFP signal was detected when GFPβ1-10-Moesin and GFPβ11-Kindlin-2 were co-transfected into EA.hy926 cells, confirming that Moesin interacts with Kindlin-2 in ECs (**Fig. 3F and G**).

To investigate whether the interaction between Kindlin-2 and Moesin also occurs in vivo and whether it changes during development, we examined the retinal and brain vasculature during key stages of vascular development. In the retina, the superficial vascular plexus forms between postnatal days P1-P7, while the development of the deep retinal vasculature begins at P7 and completes by P14. PLA assays performed on retinal sections at P6, P14, and in adulthood revealed that Kindlin-2–Moesin interactions were present at all stages, with the strongest signal detected at P6, a highly angiogenic stage (**Fig. 3H and I**). In the brain, vascularization of the neural tube begins around E8.5–E10, with vessel sprouts emerging from the perineural vascular plexus to invade the central nervous system (*34*). PLA assays conducted on brain sections from mouse embryos at E11.5, E12.5, E13.5, and E14.5 showed that the Kindlin-2-Moesin interaction was significantly higher at the early, highly angiogenic stages (E11.5 and E12.5) compared to the later, less angiogenic stages (E13.5 and E14.5) (**Fig. 3J and K**).

### Kindlin-2 interacts with N62 residue of Moesin to regulate Moesin activity and membrane tension

To further map the binding region of Kindlin-2 and Moesin, we generated three fragments of Kindlin-2—N-terminus (NT), middle region (MR), and C-terminus (CT)—each tagged with Flag, and co-transfected them with HA-tagged Moesin into HEK293T cells. Co-immunoprecipitation (Co-IP) analysis using anti-Flag beads revealed that the NT of Kindlin-2 specifically interacted with HA-tagged Moesin, whereas the MR and CT fragments showed no detectable binding (**Fig. 4A**). Similarly, we generated two Moesin fragments (NT and CT) tagged with HA and co-transfected them into HEK293T cells with Flag-tagged Kindlin-2. Anti-Flag Co-IP demonstrated that the NT of Moesin interacted with Flag-tagged Kindlin-2, while the CT of Moesin did not (**Fig. 4B**). Together, these results indicate that the N-terminal regions of both Kindlin-2 and Moesin mediate their interaction.

**Figure 4.**
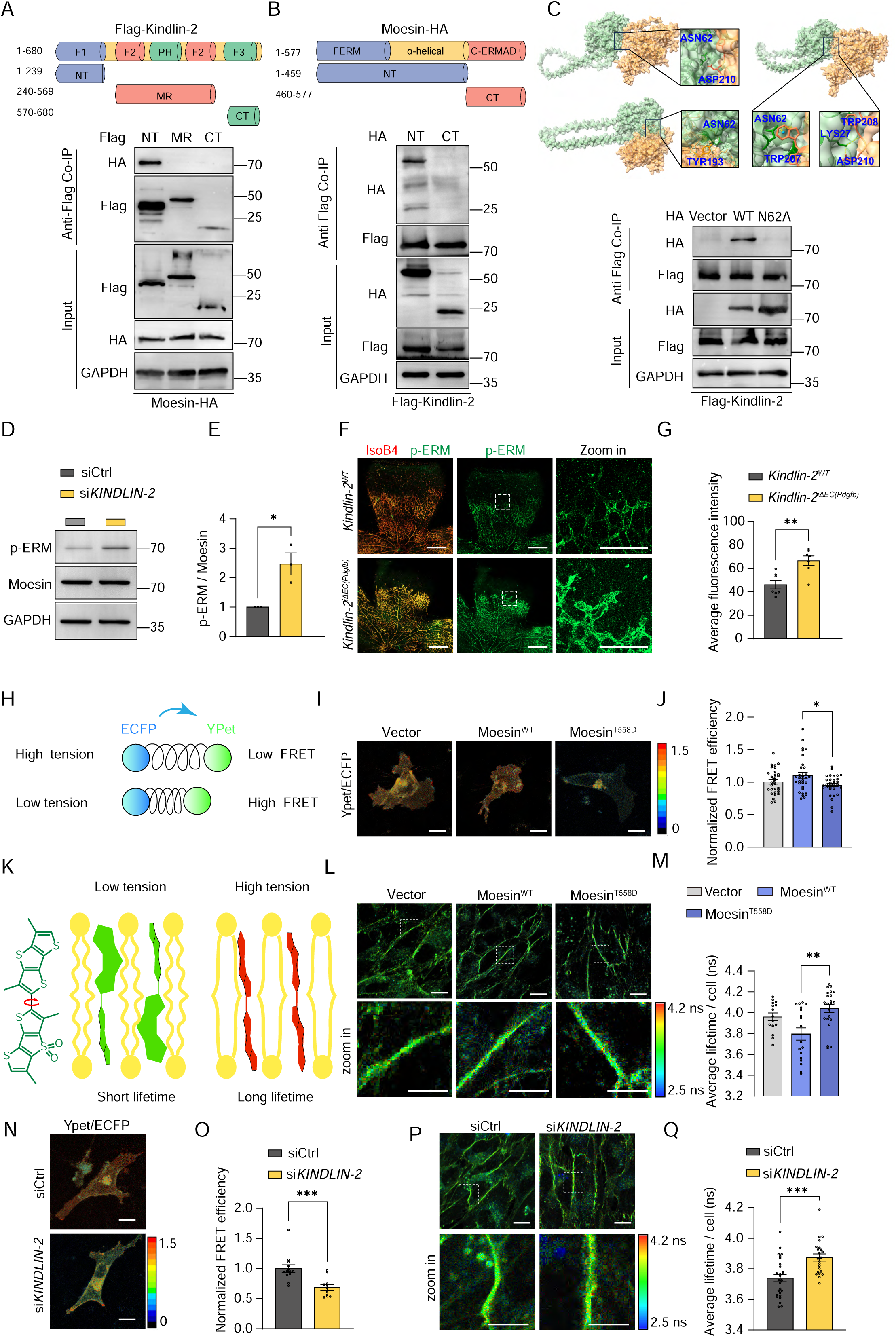
Kindlin-2 interacts with Moesin at N62 to regulate Moesin activity and membrane tension. (A) Schematic diagram of three truncation mutants of Kindlin-2. Western blot analysis of HEK293T cells co-transfected with Flag-tagged NT, MR, or CT fragments of Kindlin-2 and HA-tagged Moesin for 48 hrs, followed by immunoprecipitation with anti-Flag M2 magnetic beads. Blots shown are representative of 3 independent experiments. (B) Schematic diagram of two truncation mutants of Moesin. Western blot analysis of HEK293T cells co-transfected with HA-tagged NT or CT fragments of Moesin and Flag-Kindlin-2, followed by immunoprecipitation with anti-Flag M2 beads. Blots are representative of 3 independent experiments. (C) Structural modeling of the Kindlin-2–Moesin interface. Kindlin-2 and Moesin residues are shown in green and yellow, respectively. Structure predictions were generated using AlphaFold and ChimeraX. Moesin residue N62 was predicted to be critical for Kindlin-2 binding in three models. To validate this, HEK293T cells were transfected with HA-Moesin^WT^ or HA-Moesin^N62A^ and analyzed for Kindlin-2 binding via Co-IP. Blots are representative of 3 independent experiments. (D) Western blot analysis of HUVECs transfected with control or *KINDLIN-2* siRNA for 48 hrs. (E) Quantification of p-ERM levels from (D) (n = 3 independent experiments). (F) Representative flat-mounted retinal images from P6 *Kindlin-2^WT^*and *Kindlin-2^iΔEC(Pdgfb)^* pups stained for p-ERM (green) and IsoB4 (red), highlighting elevated p-ERM levels at the angiogenic front of *Kindlin-*2*^iΔEC(Pdgfb)^* pups. (G) Quantification of p-ERM fluorescence intensity in IsoB4^+^ vascular regions from (F) (n = 7/7 pups). (H) Schematic of the membrane tension biosensor MSS, a membrane-bound FRET-based probe consisting of an elastic spider silk domain between ECFP and YPet. Membrane tension changes are detected through changes in FRET efficiency. (I) Representative ratiometric FRET images of MSS probe in EA.hy926 cells infected with Ad-Vector, Ad-Moesin^WT^, or Ad-Moesin^T558D^ (constitutively active Moesin) for 48 hrs. (J) Quantification of FRET efficiency in (I) (n = 31/31/32 cells from 3 independent experiments). (K) Schematic of the Flipper-TR probe, which integrates into the plasma membrane and reports tension changes through shifts in fluorescence lifetime. (L) Representative fluorescence lifetime images of HUVECs infected with Ad-Vector, Ad-Moesin^WT^, or Ad-Moesin^T558D^. Cells were incubated with 1.5 μM Flipper-TR for 15 mins at 37 °C before fluorescence lifetime imaging microscopy (FLIM) imaging. (M) Quantification of fluorescence lifetime from (L) (n = 15/18/22 cells from 3 independent experiments). (N) Representative ratiometric FRET images of EA.hy926 cells transfected with siCtrl or si*KINDLIN-2*. (O) Quantification of FRET efficiency from (N) (n = 13/11 cells from 3 independent experiments). (P) Representative fluorescence lifetime images of HUVECs transfected with siCtrl or si*KINDLIN-2*. (Q) Quantification of fluorescence lifetime from (P) (n = 27/23 cells from 3 independent experiments). Data are presented as mean ± SEM. * *P* < 0.05; ** *P* < 0.01; *** *P* < 0.001 by two-tailed Student’s t-test or one-way ANOVA followed by Tukey’s multiple comparisons test. Scale bars: 500 μm in low magnification of (F); 100 μm in high magnification of (B) and (E); 5 μm in (I), (N), and low magnification of (L) and (P); 2.5 μm in high magnification of (L) and (P).

To further explore the structural basis of the Kindlin-2-Moesin interaction, we used AlphaFold to predict their potential interaction interfaces. Among the multiple binding modes predicted by AlphaFold, Moesin N62 residue was consistently identified as a key interface for binding with Kindlin-2 (**Fig. 4C**). Based on this, we generated a point mutation, substituting Moesin N62 with alanine. Co-IP analysis revealed that Moesin^N62A^ completely abolished the binding with Kindlin-2 (**Fig. 4C**), indicating that Kindlin-2 binds to Moesin through Moesin N62 residue.

We next investigated the functional consequences of the interaction between Kindlin-2 and Moesin. The ERM protein family link cortical cytoskeleton to the plasma membrane through a C-terminal F-actin binding domain and an N-terminal membrane-binding module (*35*). In their inactive state, ERM proteins adopt a closed conformation in the cytosol, maintained by intramolecular interactions between their N- and C-terminal domains, which mask the F-actin binding site. Activation of ERM proteins requires two sequential regulatory events: binding of phosphatidylinositol 4,5-bisphosphate (PIP2) to the N-terminal domain, followed by phosphorylation of a conserved C-terminal threonine residue (T567 in Ezrin, T564 in Radixin, and T558 in Moesin) (*36, 37*). Notably, lysine 63 (K63) in Moesin is recognized as a crucial residue for PIP2 binding (*38*). Based on this, we hypothesized that Kindlin-2 binding to Moesin at Moesin^N62^ may interfere with PIP2-Moesin interaction, thereby limiting Moesin phosphorylation and activation. Conversely, loss of Kindlin-2 may lead to constant PIP2-Moesin interaction, resulting in hyper phosphorylation of Moesin (**Fig. S4B**).

Indeed, upon knocking down Kindlin-2 in HUVECs, we observed a significant increase in phosphorylated ERM (pERM) levels (**Fig. 4D-E**). Gene expression analysis showed that Moesin expression is substantially higher than that of Radixin or Ezrin across different endothelial populations (**Fig. S4C**). Additionally, RNA-seq analysis of isolated brain ECs and whole brain tissue from mouse embryos at various developmental stages revealed that the *Msn* is significantly enriched in ECs compared to the whole tissue (**Fig. S4D**). These data suggest that the observed increase in pERM is primarily due to enhanced Moesin phosphorylation rather than Radixin or Ezrin. Consistently, in vivo analysis also revealed that pERM levels were significantly elevated in the retinal vasculature of *Kindlin-2^iΔEC(Pdgfb)^* pups at P6 (**Fig. 4F and G**).

As ERM proteins are key regulators of membrane-cortex interactions and provide resistance to the expansion of the membrane for regulating membrane tension (*39*), we next investigated whether membrane tension was altered following Kindlin-2 knockdown. To assess this, we employed two complementary methods: the membrane stress sensor (MSS) biosensor and the fluorescent lipid tension reporter Flipper-TR. MSS is a membrane-bound fluorescence resonance energy transfer (FRET)-based sensor in which FRET efficiency decreases under high membrane tension and increases when membrane tension is low (*40, 41*). Flipper-TR is a fluorescent probe that integrates into the plasma membrane and reports membrane tension by exhibiting longer fluorescence lifetimes under increased tension and shorter lifetimes under reduced tension (*41*). To validate the system, we generated an adenovirus expressing a constitutively active form of Moesin (Moesin^T558D^), which mimics Thr558-phosphorylated state of Moesin. Infection of HUVECs with Ad-Moesin^T558D^ significantly reduced FRET efficiency in the MSS assay and prolonged the fluorescence lifetime of Flipper-TR, indicating increased membrane tension (**Fig. 4H-M**). Similarly, *KINDLIN-2* knockdown also resulted in increased membrane tension, as detected by both methods (**Fig. 4N-Q**).

### Kindlin-2/Moesin interaction regulates VEGFR2 endocytosis and VEGF signaling

To understand the biological consequences of Moesin hyperactivation, we infected HUVECs with Ad-Moesin^T558D^ or control wild type Ad-Moesin in HUVECs and performed RNA sequencing. 1084 genes were upregulated, while 1117 genes were downregulated upon Moesin^T558D^ expression. Pathway enrichment analysis of the altered transcripts revealed that the most significant changed pathway in the VEGFA-VEGFR2 signaling pathway (**Fig. 5A**).

**Figure 5.**
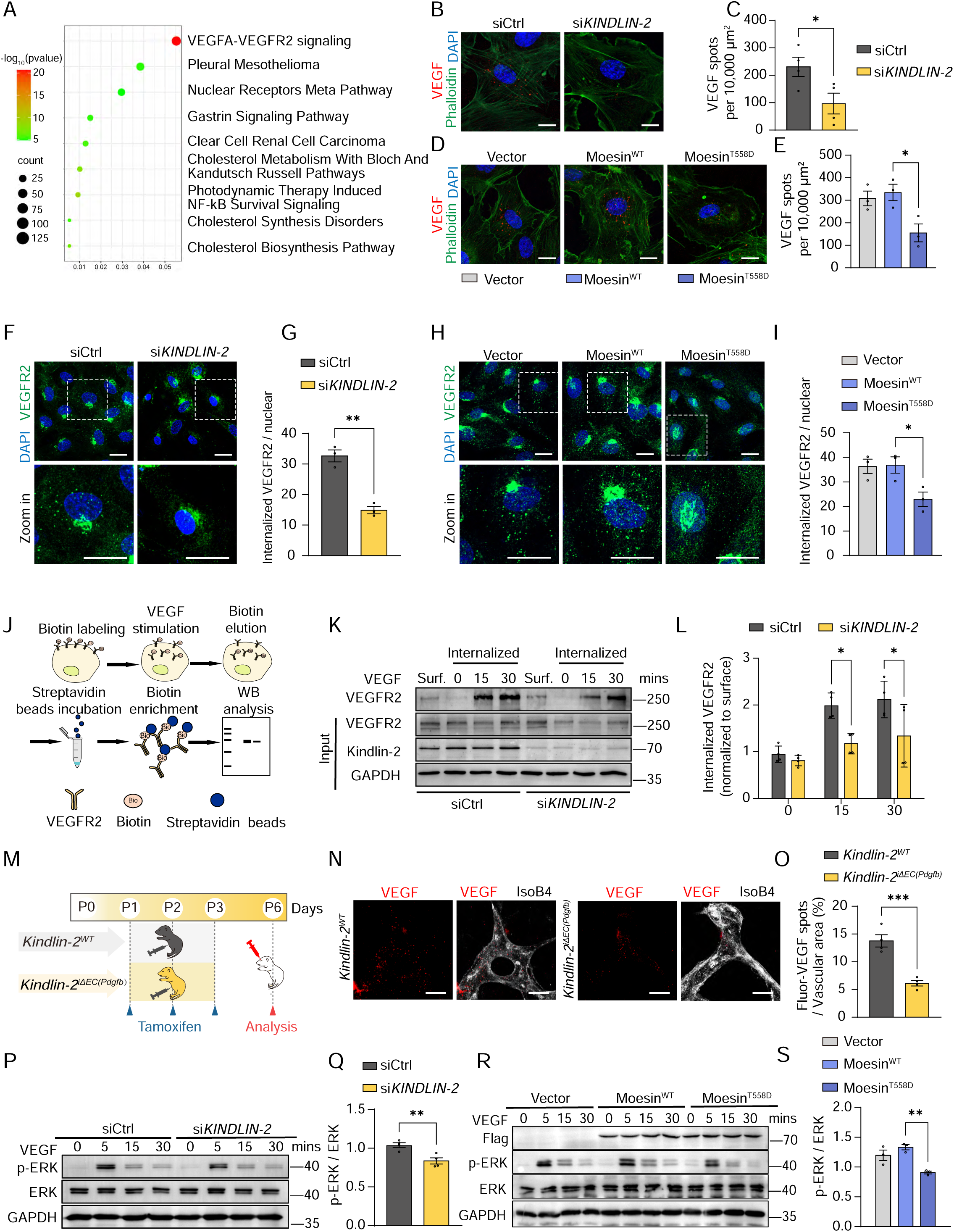
Kindlin-2/Moesin interaction regulates VEGFR2 endocytosis and VEGF signaling. (A) HUVECs were infected with Ad-Moesin^WT^ or Ad-Moesin^T558D^ for 48 hrs, and total RNA was harvested for RNA-seq. The differentially expressed genes were analyzed with Gene Ontology Enrichment Analysis. The top nine gene sets are shown. Data were analyzed from GSE302986. (B) HUVECs transfected with siCtrl or si*KINDLIN-2* were serum-starved (2% FBS) for 8 hrs, then stimulated with 2 μg/mL Alexa 594-labeled VEGF (red) for 30 mins. VEGF accumulation was visualized by confocal microscopy. Actin, phalloidin (green); nuclei, DAPI (blue). (C) Quantification of Alexa 594-labeled VEGF spots normalized to cellular area (n= 4 independent experiments). (D) HUVECs infected with Ad-Moesin^WT^ or Ad-Moesin^T558D^ were starved and stimulated with Alexa 594-labeled VEGF as in (B). VEGF accumulation was imaged by confocal microscopy. (E) Quantification of Alexa 594-labeled VEGF spots normalized to cellular area (n=3 independent experiments). (F) HUVECs transfected with siCtrl or si*KINDLIN-2* were serum-starved for 8 hrs, stimulated with 50 ng/ml VEGF for 30 mins, fixed, and stained for VEGFR2 (green). Representative images show intracellular VEGFR2 vesicles. (G) Quantification of VEGFR2 vesicle number per cell from (F) (n=3 independent experiments). (H) HUVECs infected with Ad-Moesin^WT^ or Ad-Moesin^T558D^ were treated as in (F), and stained for VEGFR2. (I) Quantification of VEGFR2 vesicle number per cell from (H) (n= 3 independent experiments). (J) Schematic of the cell surface biotinylation assay. HUVECs were starved for 8 hrs, labeled with EZ-Link Sulfo-NHS-SS-Biotin (0.25 mg/mL) at 4 °C for 1 hr, then stimulated with VEGF (50 ng/mL) for 30 mins. After surface biotin was stripped with GSH elution buffer, total proteins were extracted. Biotinylated internalized VEGFR2 was pulled down using streptavidin magnetic beads and analyzed by western blotting. (K) Cell surface biotinylation assay for VEGFR2 internalization in siCtrl- and si*KINDLIN-2*–transfected HUVECs. Input lysates show VEGFR2, Kindlin-2, and GAPDH. “Surf” represents surface VEGFR2 prior to VEGF stimulation and biotin stripping. (L) Quantification of internalized VEGFR2 normalized to surface VEGFR2 levels (n=4 independent experiments). (M) Schematic of in vivo Alexa 594-labeled VEGF uptake assay in *Kindlin-2^WT^* and *Kindlin-2^iΔEC(Pdgfb)^* pups. Tamoxifen was administered from P1 to P3; Alexa 594-labeled VEGF was injected intravitreally at P6 and analyzed after 30 mins. (N) Representative confocal images of the retinal angiogenic front showing uptake of Alexa 594-labeled VEGF by ECs from *Kindlin-2^WT^*and *Kindlin-2^iΔEC(Pdgfb)^* pups. (O) Quantification of internalized Alexa 594-VEGF at the angiogenic front (n = 4/4 pups). (P) Western blot of HUVECs transfected with siCtrl or si*KINDLIN-2*, starved for 8 hrs, then stimulated with 50 ng/mL VEGF. (Q) Quantification of p-ERK levels at 5 mins after VEGF stimulation from (P) (n = 4 independent experiments). (R) Western blot of HUVECs infected with Ad-Vector, Ad-Moesin^WT^, or Ad-Moesin^T558D^, treated as in (P). (S) Quantification of p-ERK levels at 5 mins after VEGF stimulation from (R) (n = 3 independent experiments). Data are presented as mean ± SEM. * *P* < 0.05; ** *P* < 0.01; *** *P* < 0.001 by two-tailed Student’s t-test or one-way ANOVA followed by Tukey’s multiple comparisons test. Scale bars: 5 μm in (B), (D), (F), (H), and (N).

VEGFR2, like other receptor tyrosine kinases (RTKs), is known to signal from the plasma membrane following ligand-induced dimerization and activation. Its endocytosis and subsequent trafficking are key steps in regulating its signaling (*42*). Notably, endocytosis is a process that is known to be influenced by membrane tension (*43–45*). Therefore, we investigated whether Moesin^T558D^ expression or *KINDLIN-2* knockdown would affect VEGFR2 endocytosis.

To visualize endocytosis, we used Alexa 594-labeled human VEGF to stimulate HUVECs transfected with siCtrl or si*KINDLIN-2*, or infected with Ad-Vector, Ad-Moesin^WT^ or Ad-Moesin^T558D^. Both si*KINDLIN-2* and Moesin^T558D^ expression resulted in reduced accumulation of VEGF in the cells (**Fig. 5B-E**). Immunofluorescence staining of VEGFR2 further revealed a marked decrease in intracellular VEGFR2 in both the si*KINDLIN-2* and Moesin^T558D^ groups, indicating impaired endocytosis (**Fig. 5F-I**). To validate this finding, we performed a cell surface biotinylation assay (*46*), which confirmed that *KINDLIN-2* knockdown significantly reduced VEGF-induced VEGFR2 endocytosis (**Fig. 5J-L**). To assess VEGF uptake in vivo, Alexa dye–conjugated VEGF was injected intravitreally into P6 pups as previous described (*47*)(**Fig. 5M**). The accumulation of labeled VEGF could be observed in retinal ECs at the angiogenic front 30 minutes post-injection. Notably, the Fluor-VEGF spots were significantly reduced in *Kindlin-2^iΔEC(Pdgfb)^* mice compared with *Kindlin-2^WT^*controls (**Fig. 5N and O**). As ERK activation is a well-established endocytosis-dependent downstream response of VEGFR2 signaling (*42*), we examined ERK phosphorylation. Consistent with impaired receptor internalization, VEGF-induced ERK activation was reduced in both the si*KINDLIN-2* and Moesin^T558D^ groups (**Fig. 5P-S**).

To further confirm that membrane tension regulates VEGFR2 endocytosis, we used an established cell culture model: hypotonic treatment, known to increase membrane tension (*48*). For this treatment, HUVECs were exposed to hypotonic medium, as previously described (*48*). MSS biosensor analysis showed a reduction in FRET efficiency, confirming successful induction of membrane tension (**Fig. S5A and B**). Alexa dye-coupled VEGF feeding assay further demonstrated that hypotonic exposure significantly decreased VEGF uptake (**Fig. S5C and D**). Consistently, p-ERK levels were also reduced in HUVECs exposed to the hypotonic treatment following VEGF stimulation (**Fig. S5E and F**).

### Deficiency of Kindlin-2 impairs VEGF signaling in a Moesin-dependent manner

To determine whether the impaired VEGF signaling and angiogenesis observed in Kindlin-2-deficient cells are Moesin-dependent, we co-knocked down *MSN* in Kindlin-2-deficient cells to assess potential rescue effects. We found that *KINDLIN-2* knockdown significantly decreased MSS FRET efficiency, whereas co-transfection with *MSN* siRNA effectively restored FRET efficiency (**Fig. 6A and B**). Similarly, the fluorescence lifetime of Flipper-TR, which increased in Kindlin-2-deficient ECs, was reduced upon *MSN* knockdown (**Fig. S6A and B**).

**Figure 6.**
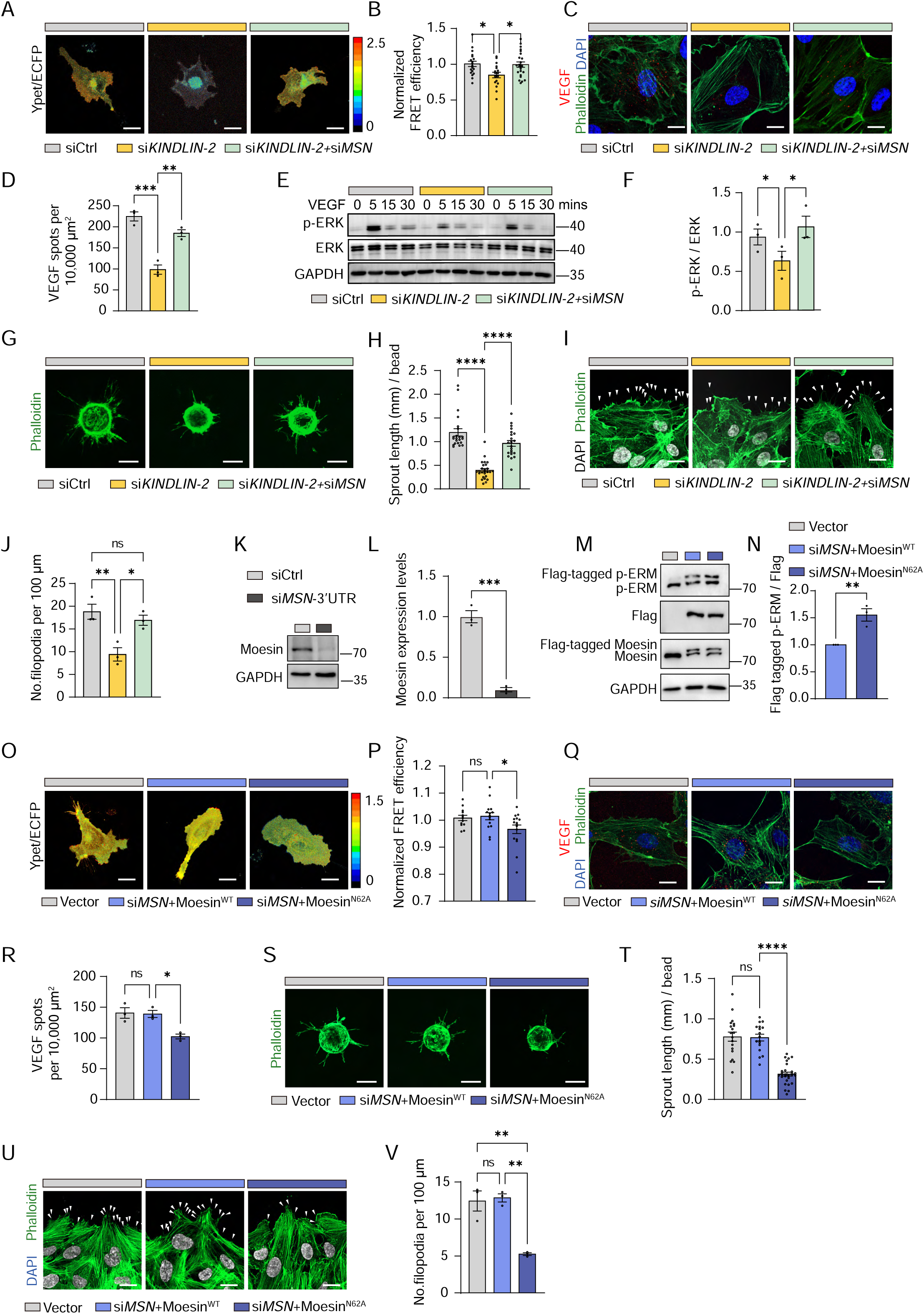
Deficiency of Kindlin-2 impairs VEGF signaling in a Moesin-dependent manner. (A) Representative FRET ratiometric images of the MSS probe in EA.hy926 cells transfected with siCtrl, si*KINDLIN-2*, or si*KINDLIN-2* + si*MSN* for 48 hours. (B) Quantification of FRET efficiency in (A) (n = 19/17/23 cells from 3 independent experiments). (C) HUVECs transfected with siCtrl, si*KINDLIN-2*, or si*KINDLIN-2* + si*MSN* were serum-starved (2% FBS) for 8 hrs and stimulated with 2 μg/mL Alexa 594-labeled VEGF (red) for 30 mins. VEGF accumulation was visualized by confocal microscopy. Actin, phalloidin (green); nuclei, DAPI (blue). (D) Quantification of intracellular Alexa 594-labeled VEGF spots (n=3 independent experiments). (E) HUVECs were treated as in (C) and stimulated with 50 ng/mL VEGF. Western blot analysis shows ERK phosphorylation 5 mins after VEGF stimulation. (F) Quantification of p-ERK levels at 5 mins after VEGF stimulation in (E) (n = 3 independent experiments). (G) Representative images from the bead-sprouting assay using HUVECs transfected with siCtrl, si*KINDLIN-2*, or si*KINDLIN-2* + si*MSN* and cultured for 48 hrs. Actin, phalloidin (green). (H) Quantification of total sprout length per bead in (G) (n = 22/26/22 beads from 3 independent experiments). (I) HUVECs were transfected with siCtrl, si*KINDLIN-2*, or si*KINDLIN-2* + si*MSN* for 48 hrs, 8 hours serum-starved (2% FBS), and scratched. After VEGF treatment (50 ng/mL) for 6 hrs, cells were fixed and stained with phalloidin. White arrows indicate phalloidin⁺ filopodia. (J) Quantification of phalloidin⁺ filopodia from (I) (n = 3 independent experiments). (K) Western blot analysis of Moesin expression after si*MSN*-3’UTR transfection in HUVECs for 48 hrs. (L) Quantification of knockdown efficiency from (K). (M) HUVECs were transfected with siCtrl or si*MSN*-3’UTR and infected with Ad-Vector, Ad-Moesin^WT^, or Ad-Moesin^N62A^ for 48 hrs. Moesin phosphorylation was assessed by western blotting. Lower band indicates endogenous Moesin and p-ERM; upper band represents exogenous Flag tagged Moesin or Flag tagged p-ERM. (N) Quantification of p-ERM levels from (M) (n = 3 independent experiments). (O) Representative FRET ratiometric images of the MSS probe in EA.hy926 cells transfected with siCtrl or si*MSN*-3’UTR and infected with Ad-Vector, Ad-Moesin^WT^, or Ad-Moesin^N62A^ for 48 hrs. (P) Quantification of FRET efficiency in (M) (n = 13/16/28 cells from 3 independent experiments). (Q) HUVECs were transfected with siCtrl or si*MSN*-3’UTR and infected with Ad-Vector, Ad-Moesin^WT^, or Ad-Moesin^N62A^ for 48 hrs, then serum-starved (2% FBS) for 8 hrs and stimulated with 2 μg/mL Alexa 594-labeled VEGF for 30 mins. VEGF accumulation was visualized by confocal microscopy. Actin, phalloidin (green); nuclei, DAPI (blue). (R) Quantification of intracellular Alexa 594-labeled VEGF spots from (O) (n=3 independent experiments). (S) Representative images from the bead-sprouting assay using HUVECs transfected with siCtrl or si*MSN*-3’UTR and infected with Ad-Vector, Ad-Moesin^WT^, or Ad-Moesin^N62A^, cultured for 48 hrs. Actin, phalloidin (green). (T) Quantification of total sprout length per bead in (S) (n = 20/17/28 beads from 3 independent experiments). (U) HUVECs were transfected and treated as in (S). After scratch wounding and VEGF (50 ng/mL) treatment for 6 hrs, cells were fixed and stained with phalloidin. White arrows indicate phalloidin⁺ filopodia. Actin, phalloidin (green), nuclei, DAPI (White). (V) Quantification of phalloidin⁺ filopodia from (U) (n = 3 independent experiments). Data are presented as mean ± SEM. * *P* < 0.05; ** *P* < 0.01; *** *P* < 0.001; **** *P* < 0.0001 by one-way ANOVA followed by Tukey’s multiple comparisons test. Scale bars: 100 μm in (G) and (S); 5 μm in (A), (C), (I), (O), (Q), and (U).

VEGF feeding assays and immunofluorescence staining revealed that VEGFR2 endocytosis was reduced in si*KINDLIN-2* ECs compared to siCtrl ECs, whereas co-knockdown of *MSN* in si*KINDLIN-2* ECs significantly rescued this defect (**Fig. 6C and D**, **Fig. S6C and D**). Moreover, si*MSN* also restored the reduction in p-ERK levels caused by *KINDLIN-2* knockdown in response to VEGF stimulation (**Fig. 6E and F**).

To analyze the functional consequences, we performed in vitro bead sprouting and filopodia formation assays. The results showed that transfection of si*MSN* successfully rescued the defects in sprouting and filopodia formation caused by *KINDLIN-2* knockdown (**Fig. 6G-J**).

As we have proved N62 on Moesin is a critical site for the interaction with Kindlin-2, we constructed an adenovirus expressing Flag-tagged Moesin^N62A^ (Ad-Moesin^N62A^), a mutant unable to bind to Kindlin-2. We knocked down endogenous Moesin in HUVECs using siRNA targeting the 3’ UTR of *MSN* and then infected the cells with either Ad-Flag-Moesin^N62A^ or the control Ad-Flag-Moesin. Knockdown efficiency of si*MSN*-3’UTR was confirmed by western blotting (**Fig. 6K and L**). We found that Ad-Moesin^N62A^ phenocopied the effects of *KINDLIN-2* knockdown, resulting in increased p-ERM levels and membrane tension, along with decreased VEGF uptake and VEGFR2 endocytosis (**Fig. 6M-R**, **Fig. S6E and F**). Consequently, this led to defects in sprouting and filopodia formation (**Fig. 6S-V**).

### Kindlin-2–Moesin interaction is involved in pathological angiogenesis

Given that Kindlin-2 inhibition reduced VEGFR2 endocytosis and suppressed angiogenesis, we next sought to investigate whether Kindlin-2 also contributes to pathological angiogenesis. To this end, we first utilized the oxygen-induced retinopathy (OIR) model, which mimics the pathological features of proliferative diabetic retinopathy (PDR) and retinopathy of prematurity (ROP) (**Fig. 7A**). In this model, we exposed pups to 75% oxygen to induce vessel regression from P7 to P12. Subsequently, the pups were returned to normoxia until P17, and the relative hypoxia triggers VEGF upregulation, which drives pathological vessel sprouting and the formation of abnormal vascular tufts. To assess Kindlin-2 involvement in OIR, we analyzed single-cell RNA-seq data from P17 OIR retinas (GSE216676). Kindlin-2 was highly expressed in both tuft and tip ECs, suggesting a role in tuft formation and vascularization (**Fig. S7A and B**). PLA further indicated that the interaction between Kindlin-2 and Moesin is increased in the vasculature of the OIR retina, implying that their interaction is particularly pronounced in highly angiogenic blood vessels (**Fig. 7B and C**).

**Figure 7.**
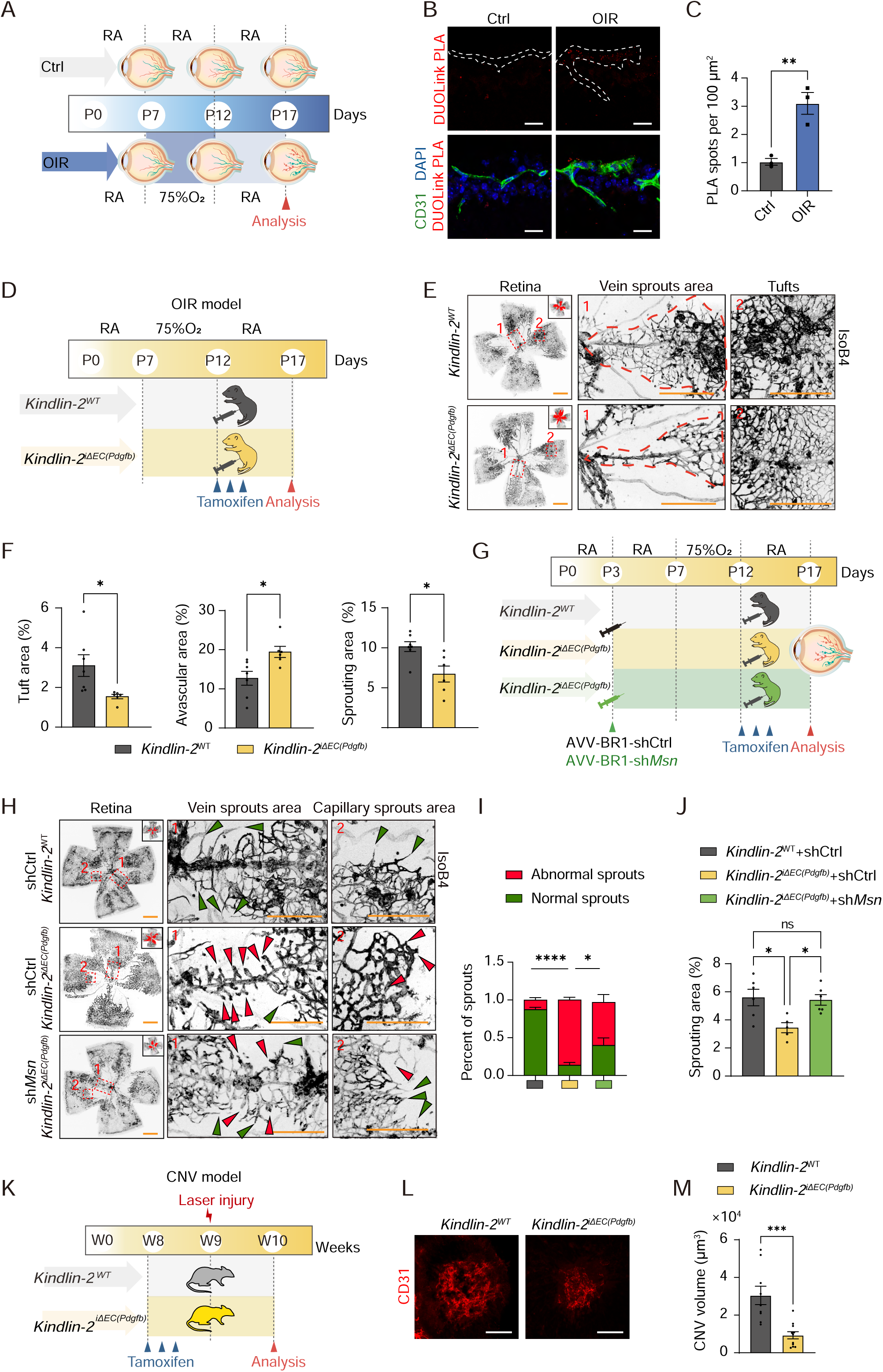
Kindlin-2–Moesin interaction is involved in pathological angiogenesis. (A) Schematic diagram of the oxygen-induced retinopathy (OIR) model. Pups were exposed to 75% oxygen from P7 to P12 and then returned to room air (RA) until P17. (B) Representative images from proximity ligation assay (PLA) for Kindlin-2 and Moesin on sagittal cryosections of eyes from control and OIR pups. Blood vessels were stained with CD31 (green), nuclei with DAPI (blue), and in situ PLA signals (red) indicate Kindlin-2–Moesin interactions. White dotted line indicates the CD31^+^ area. (C) Quantification of PLA signal spots in CD31⁺ vascular areas from (B) (n = 3/3/3 pups). (D) Schematic of Tamoxifen injection in the OIR model. *Kindlin-2^WT^*and *Kindlin-2^iΔEC(Pdgfb)^* pups were exposed to 75% oxygen from P7 to P12, returned to room air, and injected with Tamoxifen at P12, P13, and P14. Retinas were collected for analysis at P17. (E) Representative images of IsoB4-stained retinal vasculature from *Kindlin-2^WT^* and *Kindlin-2^iΔEC(Pdgfb)^* pups at P17 in the OIR model. (F) Quantification of neovascular tuft area, avascular area, and vein sprouting area in *Kindlin-2^WT^* and *Kindlin-2^iΔEC(Pdgfb)^* retinas (n = 7/6 pups). (G) Schematic of AAV-BR1 injection in the OIR model. AAV-shCtrl or AAV-sh*Msn* was administrated via retro-orbital injection into *Kindlin-2^WT^* and *Kindlin-2^iΔEC(Pdgfb)^* pups at P3. Pups were exposed to 75% oxygen from P7 to P12, then returned to room air. Tamoxifen was administered at P12, P13, and P14. Retinas were analyzed at P17. (H) Representative images of IsoB4-stained retinal vasculature from *Kindlin-2^WT^* and *Kindlin-2^iΔEC(Pdgfb)^* pups injected with AAV-shCtrl or AAV-sh*Msn*. Green arrows indicate normal sprouts from veins or capillaries; red arrows indicate blunt sprouts. (I) Quantification of the percentage of abnormal versus normal sprouts from veins in (H) (n=6/5/6 pups). (J) Quantification of the sprouting area from veins in (H) (n=6/5/6 pups). (K) Schematic of the choroidal neovascularization (CNV) model. Eight-week-old mice received 3 Tamoxifen injections every other day. CNV was induced two days after the final injection. (L) Representative confocal images of CD31 (red)-stained RPE–choroid–sclera flat mounts from CNV mice 7 days post-laser injury. (M) Quantification of CNV volume, calculated as total CD31⁺ volume at the laser lesion site (n = 9/11 mice). Data are shown as mean ± SEM. * *P* < 0.05; ** *P* < 0.01; *** *P* < 0.001, one-way ANOVA followed by Tukey’s multiple comparisons test. Scale bars: 500 μm in low-magnification panels of (D) and (H); 200 μm in high-magnification panels of (D), (E), and (I); 100 μm in (K); 5 μm in (B).

To evaluate the effect of Kindlin-2 loss on pathological retinal neovascularization, Cre recombination was induced after the return to normoxia (P12 to P14) (**Fig. 7D**). Analysis at P17 revealed that pathological neovascularization, quantified by the neovascular tuft area, was significantly reduced in *Kindlin-2^iΔEC(Pdgfb)^* mice compared with littermate controls (**Fig. 7E and F**). Additionally, assessment of the avascular area showed impaired revascularization (**Fig. 7F**). Furthermore, the sprouting area in the avascular region was also reduced (**Fig. 7F**).

Next, we aimed to investigate whether the defects observed in *Kindlin-2*-deficient mice could be rescued by Moesin knockdown. To this end, we used an AAV-BR1 system, which allows for specific gene delivery to the CNS vasculature including the retina (**Fig. S7C**), as previously described (*49, 50*). We constructed an AAV-BR1 vector expressing shRNA targeting *Msn* in mice and successfully reduced *Msn* expression following retro-orbital injection (**Fig. S7D**). We then injected AAV-BR1-shCtrl and AAV-BR1-*shMsn* into *Kindlin-2^iΔEC(Pdgfb)^*pups, while AAV-BR1-shCtrl-injected *Kindlin-2^WT^* pups served as controls (**Fig. 7G**). Similar to observations in the P6 retina, tip cells in the avascular area of the *Kindlin-2^iΔEC(Pdgfb)^* retina displayed a blunted-end, and knockdown of *Msn* in *Kindlin-2^iΔEC(Pdgfb)^*pups successfully rescued the tip cell defects caused by *Kindlin-2* loss (**Fig. 7H and I**). Consequently, the sprouting area in the avascular region was also rescued by *Msn* knockdown (**Fig. 7J**).

As the OIR model is a developmental mouse model, we next investigated whether endothelial Kindlin-2 plays a role in pathological angiogenesis during adulthood. To address this, we used a laser-induced choroidal neovascularization (CNV) mouse model, which mimics human neovascular age-related macular degeneration (nvAMD), the leading cause of blindness in the aging population (*51*). To investigate the role of Kindlin-2 in CNV formation, we injected Tamoxifen in 8-week-old *Kindlin-2^iΔEC(Pdgfb)^* mice and performed laser photocoagulation one week later (**Fig. 7K**). *Kindlin-2^iΔEC(Pdgfb)^*mice exhibited significantly reduced CNV volume compared to *Kindlin-2^WT^* mice at 7 days post-laser photocoagulation (**Fig. 7L and M**), indicating that Kindlin-2 also plays a substantial role in pathological angiogenesis during adulthood.

## Discussion

Changes in membrane mechanics are closely linked to signaling pathways and cellular fate, yet their impact on angiogenesis remains largely unexplored. In this study, we identified the interaction between Kindlin-2 and Moesin as a critical regulator of sprouting angiogenesis by modulating membrane tension. Mechanistically, we found that Kindlin-2 limits Moesin overactivation, ensuring optimal membrane tension for VEGFR2 endocytosis and consequently effective VEGF-VEGFR2 signaling. These findings establish an important link between membrane mechanics and EC behavior in both developmental and pathological angiogenesis.

Here, we highlight Kindlin-2 as a key regulator of membrane mechanics through its interaction with Moesin, a well-established mediator of membrane tension. Notably, we observed that the Kindlin-2–Moesin interaction is enhanced during highly angiogenic developmental stages, suggesting that this interaction is dynamically regulated. Interestingly, Kindlin-2 is not only a regulator for cell intrinsic mechanics, but is also recognized as a mechanoresponsive protein capable of responding to external mechanical cues. Recent studies have shown Kindlin-2 could be regulated by tension, ECM stiffness, and shear stress (*13, 17, 18*). Although this study did not aimed to identify the upstream signaling that controls the dynamic of Kindlin-2 and Moesin interaction, it is important to notice that organ development also involves extensive mechanical signature changes (e.g. fluid shear, ECM stiffness, tension, hydrostatic pressure), and mechanical forces are known to play a crucial role in organ development, morphogenesis, and tissue patterning (*52*). Therefore, it would be interesting to explore how these forces modulate the Kindlin-2-Moesin interaction and, in turn, regulate angiogenesis.

Previous studies have highlighted the link between mechanical changes and endocytosis, especially in developmental processes and stem cell differentiation (*53, 54*). In our study, we demonstrated that changes in membrane mechanics significantly alter VEGF signaling in ECs. VEGF and their receptors play a crucial role in vascular system development. VEGFs bind with high affinity to the receptor tyrosine kinases VEGFR1–R3, with VEGFR2 being the main signaling receptor in blood vascular ECs. According to the consensus model for ligand-induced activation of RTKs, VEGF binding to its cognate VEGFR induces receptor homodimerization or heterodimerization, leading to the activation of the tyrosine kinase and subsequent autophosphorylation of tyrosine residues within the receptor intracellular domains. This activation triggers various intracellular signaling pathways, including PI3K/AKT, MAPK, and SRC signaling, which mediate cellular processes such as survival, proliferation, and migration. Endocytosis and the subsequent receptor signaling have been identified as critical determinants of signaling output. Until now, the primary known endocytic route for VEGFR2 has been the clathrin-mediated endocytic pathway, with ERK activation occurring downstream of clathrin-mediated VEGFR2 endocytosis (*55*). Moreover, alternative clathrin-independent endocytosis routes, such as fast endophilin-mediated endocytosis, micropinocytosis, and caveolae, have also been identified (*46, 56–58*). Furthermore, in ECs, other membrane-bound receptors, such as Neuropilin and Tie2, also undergo endocytosis to exert their biological functions (*59, 60*). Therefore, it would be interesting to explore whether the Kindlin-2-Moesin interaction-mediated endocytosis is a conserved mechanism across other membrane-bound receptors.

The cell membrane functions as a barrier that controls the exchange of diffusible molecules. Membrane tension regulates several critical cellular processes, including motility, cell signaling, endocytosis, and mechanotransduction. Although our study primarily focused on the impaired VEGF signaling resulting from abnormal membrane tension, it is important to note that membrane tension also plays a key role in regulating cell migration, protrusion, and polarity, which are also crucial processes for sprouting angiogenesis (*61–64*). Consequently, the angiogenic defects observed in Kindlin-2-deficient cells may not be solely due to impaired VEGF signaling.

Excessive vessel growth is a hallmark of various diseases, including solid tumors and ocular diseases. In this study, we used two ocular diseases models—the OIR model and the CNV model—which represent PDR or ROP, as well as nvAMD. In both models, we found that depleting *Kindlin-2* suppressed neovascularization. However, since *Kindlin-2* depletion leads to impaired EC barrier function (*18*), its depletion might not be beneficial as a therapeutic approach. Thus, understanding the specific mechanisms by which Kindlin-2 promotes angiogenesis is crucial. In this study, we identified that Kindlin-2 binds to the N62 residue of Moesin, and mutation of N62 mimics the effects of Kindlin-2 inhibition in terms of VEGFR2 endocytosis and sprouting angiogenesis. Therefore, targeting this binding interface could provide a therapeutic strategy to limit excessive sprouting angiogenesis.

In summary, this study identifies the interaction between Kindlin-2 and Moesin and demonstrates how this interaction orchestrates membrane mechanics to orchestrate sprouting angiogenesis. We also highlight the critical role of Moesin N62 residue in facilitating this interaction, which is essential for regulating Moesin activity and function. These findings underscore the potential therapeutic value of targeting the Kindlin-2-Moesin axis in the treatment of neovascular diseases.

## Materials and Methods

### Cell Culture

Human Umbilical Vein Endothelial Cells (HUVECs) were cultured in Medium 199 (Gibco, 8123510) supplemented with 10% fetal bovine serum (FBS) (Gibco, 10099141C), 4 µg/mL endothelial cell growth factor (ECGF), and 1% penicillin/streptomycin (Gibco, 2585610), as described previously (*18*). EA. hy926 cells were cultured in DMEM (Gibco) containing 10% FBS (Hyclone), 100 units/mL penicillin, and 100 μg/mL streptomycin. All cells were maintained in a humidified incubator with 5% CO_2_ at 37 °C.

### Mice

All study protocols involving animals were approved by the Institutional Animal Care and Use Committee of Tianjin Medical University (TMUaMEC 2020006). Mice were housed under a 12-hrs light–dark cycle at 21–25 °C with 30–70% humidity and had unrestricted access to food and water. *Kindlin2^fl/fl^*mice (*27*) were crossed with *Pdgfb-CreERT2* (*28*), and *Cdh5-(PAC)-CreERT2* mice (*65*) to generate the *Kindlin-2^iΔEC^* ^(*Pdgfb*)^ and *Kindlin-2^iΔEC^* ^(*Cdh5*)^ lines for endothelial-specific deletion of *Kindlin-2*. For analysis at P6, Cre recombination was induced by intragastric injection of 50 µL tamoxifen (1 mg/mL) daily from P1 to P3. For analysis at P9, tamoxifen was administered daily from P4 to P6 (*66*). For analysis at P21, tamoxifen was administered intragastrically from P14 to P16. For adult mice, Cre recombination was induced by three consecutive daily intraperitoneal injections of 2 mg tamoxifen.

### RNA extraction and quantitative real-time PCR analysis

Cells were collected for RNA extraction using TRIzol reagent (SIGMA, #101254514). RNA concentration was determined by spectrophotometry using a NanoDrop ND-1000 (Thermo Fisher Scientific, USA). The extracted RNA was then reverse-transcribed into complementary DNA (cDNA) using an mRNA Reverse Transcription Kit (Thermo Fisher Scientific, #K1622), following the manufacturer’s instructions. Quantitative reverse transcription PCR (qRT-PCR) was performed using SYBR™ Green Master Mix (TRANS, #AQ601) on Applied Biosystems QuantStudio 3 Real-Time PCR system (Applied Biosystems USA). The primer sequences used in this study are provided in **Table S1**. All qRT-PCR results were obtained from at least three biological replicates.

### Immunoprecipitation and immunoblotting

Co-immunoprecipitation assays were conducted as previously described (*18*). Cells were lysed using mild lysis buffer [20 mM Tris at pH 7.5, 150 mM NaCl, 5 mM EDTA, 1% NP-40, 10% glycerol, 1 × protease inhibitor cocktail, and 1 × phosphatase inhibitor (Roche)]. After centrifugation of the cell lysates for 10 mins, the resulting supernatants were used for immunoprecipitation. For the immunoprecipitation of Flag-tagged proteins, supernatants were incubated with anti-Flag M2 magnetic beads (Sigma-Aldrich, #M8823). After washing the beads, the Flag-protein complex was eluted using 3 × Flag peptide (Absin, bulk-peptide). Antibodies used for immunoblotting are listed in **Table S2**.

### IP-MS analysis

#### Immunoprecipitation

HUVECs were cultured on collagen I–coated cell culture dishes in M199 medium supplemented with 10% FBS (Gibco, 10099141C), 4 µg/mL ECGF, and 1% penicillin/streptomycin (Gibco, 2585610). HUVECs were infected with Flag-Kindlin-2 or Flag-Vector adenovirus for 48 hrs. Flag-tagged Kindlin-2 protein was pulled down using anti-Flag M2 magnetic beads (Sigma-Aldrich, #M8823). The experiment was conducted using samples pooled from three biological replicates. After gel electrophoresis and Coomassie Brilliant Blue staining, the entire Flag-Kindlin-2 and Flag-Vector gel bands were excised separately and subjected to tryptic digestion.

#### Trypsin digestion

Gel pieces were destained with 25 mM ammonium bicarbonate in ethanol/water (1:1, v/v). The destained gel pieces were then rinsed in an acidic buffer (acetic acid/ethanol/water, 1:5:4, v/v/v) twice for 1 hr each, followed by two rinses in water for 10 mins each. The gel pieces were dehydrated in acetonitrile and dried in a SpeedVac (Thermo Fisher). Disulfide bonds were reduced using 10 mM dithiothreitol at 56 °C for 1 hr and alkylated with 55 mM iodoacetamide at room temperature for 45 mins. After two water washes, the gel pieces were again dehydrated in acetonitrile and dried in a SpeedVac. Next, 200 nanograms of trypsin (Promega), dissolved in 50 mM ammonium bicarbonate, was added to the dried gel pieces and incubated overnight at 37 °C. Tryptic peptides were sequentially extracted using 50% acetonitrile (acetonitrile/water/TFA, 50:45:5, v/v/v), 75% acetonitrile (acetonitrile/water/TFA, 75:25:0.1, v/v/v), and acetonitrile. The peptide extracts were pooled, dried in a SpeedVac, and desalted using a μ-C18 Ziptip.

#### MS/MS Analysis

The digested samples were injected into a Nano-LC system (EASY-nLC 1200, Thermo Fisher Scientific). Each sample was separated using a C18 column (75 μm inner diameter × 25 cm, 3 μm C18) at a flow rate of 300 nL/min. The HPLC gradient was as follows: 5% to 10% solvent B (0.1% formic acid in 80% acetonitrile) in 16 mins, 10% to 22% solvent B in 35 mins, 22% to 30% solvent B in 15 mins, 30% to 90% solvent B in 1 min, and held at 90% solvent B for 8 mins. The HPLC eluate was electrosprayed directly into an Orbitrap Eclipse mass spectrometer (Thermo Fisher Scientific). The source operated at 2.2 kV. Mass spectrometric analysis was performed in data-dependent mode. For the MS1 survey scan, the automatic gain control (AGC) target was set to 5e4, and the resolution was 70,000. The MS2 spectra were acquired at a resolution of 17,500.

### Proximity Ligation Assay (PLA)

PLA was performed using the Duolink^®^ In Situ Red Starter Kit (Sigma, #DUO92008). For HUVECs, cells cultured on collagen-coated coverslips were fixed with 2% PFA for 10 mins at room temperature, washed with PBS, permeabilized with 0.3% Triton X-100 for 30 mins, and blocked with 0.5% Triton X-100 and 3% BSA for 1 hr at room temperature. Samples were then incubated overnight at 4 °C with anti-Moesin (Cell Signaling Technology, #3150, 1:50) and anti-Kindlin-2 (Millipore, #MAB2617, 1:50) antibodies diluted in blocking solution. Next, pre-diluted anti-rabbit PLUS and anti-mouse MINUS probes were applied for 1 hr at 37 °C, followed by ligation at 37 °C for 30 mins and amplification with polymerase for 100 mins. For mouse tissue sections (eyeball and embryo), samples were washed with PBS, permeabilized with 0.5% Triton X-100 for 2 hrs, and blocked with 0.3% Triton X-100 and 3% BSA for 1 hr at room temperature. Sections were incubated overnight at 4 °C with anti-Moesin (1:50), anti-Kindlin-2 (1:50), and anti-CD31 (Millipore, #MAB1398Z, 1:100) antibodies in blocking solution. Subsequently, pre-diluted anti-rabbit PLUS, anti-mouse MINUS probes, and goat anti-Armenian Hamster Alexa Fluor 488 (1:200, Jackson, #127-545-099) were incubated for 2 hrs at 37 °C. Ligation and polymerase amplification were performed at 37 °C for 1 hr and 2 hrs, respectively. Images were acquired using a Zeiss LSM 900 confocal microscope, and PLA signals were quantified blindly using ImageJ.

### siRNA transfection

Cells were seeded at 80% confluence and transfected with either control or gene-specific siRNA (100 nM) using Lipofectamine RNAiMAX reagent (Thermo Fisher Scientific, #13778-150) according to the manufacturer’s instructions. Fresh medium was added 6 hrs after transfection. Sequences of gene-specific siRNAs are listed in **Table S3**.

### Membrane tension measurements by fluorescent lifetime imaging (FLIM)

Membrane tension was measured using the Flipper-TR probe as previously described (*41*). Cultured cells were seeded onto confocal dishes and allowed to adhere for 12 hrs in normal growth medium, which was then replaced with imaging medium containing 1.5 µM Flipper-TR. Cells were incubated at 37 °C for 15 mins to ensure proper labeling before imaging. FLIM imaging was performed on a Leica Stellaris 8 FALCON system with 488 nm excitation and 575–625 nm emission, using a pixel dwell time > 19.58 ms. A double exponential decay model was fitted to the data to confirm a good fit (chi squared < 1.5). Counts-per-pixel and lifetime-per-pixel images were exported as 256 × 256 pixels single time-point, single Z-plane images. For quantification, membrane regions were manually identified from counts-per-pixel images, and mean lifetimes were measured by applying these ROIs to the lifetime-per-pixel images.

### Filopodia formation assay in cultured HUVECs

HUVEC filopodia formation assay was performed as previously described (*67*). Cells were seeded onto coverslips at approximately 7 × 10^4^ per well, transfected with siRNA or infected with adenovirus, and cultured in M199 medium containing 10% FBS for 48 hrs. After creating a scratch wound with a 200-μL pipette tip, cells were incubated in 10% FBS M199 for 9 hrs, followed by serum-free M199 for 12 hrs. Filopodia formation was assessed by immunofluorescence staining 6 hrs after stimulation with VEGF (50 ng/mL). For analysis, more than 15 random fields of view from at least three independent experiments were analyzed.

### In vivo VEGF internalization assay

VEGF-A (Sino Biological, #11066-HNAB) was labeled with Alexa Fluor™ 594 according to the manufacturer’s instructions (Invitrogen, #A10239). A total of 0.6 μL of labeled VEGF-A (0.2 mg/mL) was injected into the vitreous humor using glass capillary pipettes mounted on a micromanipulator (HAMILTON). After 30 minutes, the eyeballs were harvested and fixed in 4% PFA in PBS for 1 hr at 4 °C. Retinas were isolated, permeabilized overnight at 4 °C in PBS containing 1% Triton X-100, and then blocked with 0.5% Triton X-100 and 5% BSA for 3 hrs. Retinas were subsequently incubated with Alexa Fluor 488-conjugated isolectin GS-IB4 (Invitrogen, #I21411, 1:100) for 4 hrs at room temperature. Images were acquired using a confocal microscope (LSM 800, Carl Zeiss). The number of labeled VEGF-A puncta within isolectin GS-IB4-positive areas was quantified using ImageJ.

### In vitro VEGF feeding assay

For Alexa-labeled VEGF uptake in cultured cells, cells grown on coverslips were transfected with siRNA or infected with adenovirus. Cells were serum-starved for 8 hrs before incubation in serum-free growth medium containing 2 μg/mL Alexa 594-labeled VEGF-A at 37 °C for 30 mins. Cells were then fixed with 2% PFA for 10 mins at room temperature and stained with Alexa-conjugated phalloidin (Invitrogen) and DAPI. Images were acquired using a confocal microscope (Zeiss LSM 800). For analysis, more than 15 random fields of view (at least 50 cells) from at least three independent experiments were analyzed.

### In vitro VEGFR2 internalization assay

For the VEGFR2 internalization assay, cells cultured on coverslips were transfected with siRNA or infected with adenovirus. Cells were serum-starved for 8 hrs, then VEGF was added to the serum-free medium at a final concentration of 50 ng/mL and incubated at 37 °C for 30 mins. Afterwards, cells were fixed with 2% PFA for 10 mins at room temperature and stained with anti-VEGFR2 (Cell Signaling Technology, #2479S, 1:200) and DAPI. Images were acquired using a confocal microscope (Zeiss LSM 800). For analysis, more than 12 random fields of view (at least 80 cells) from at least three independent experiments were analyzed.

### Isolation of mouse brain ECs

Brain ECs were isolated using a magnetic cell separation method (Miltenyi Biotec, Germany) according to the manufacturer’s instructions. Briefly, freshly dissected mouse brains were transported in Hanks’ Balanced Salt Solution (HBSS; Gibco, #14025134), minced into small pieces, and digested with 1 mg/mL Collagenase/Dispase (Roche, #10269638001) and 60 U/mL DNase I (Roche, #4716728001) in 2% FBS endothelial cell medium at 37 °C for 6 mins. The resulting single-cell suspension was passed through a 40 μm cell strainer and washed with PBS containing 2% FBS and 2 mM EDTA. Cells were incubated with Myelin Removal Beads (Miltenyi, #130-096-733) to deplete myelin, followed by incubation with CD31 MicroBeads (Miltenyi, #130-097-418) to isolate ECs. Purified brain ECs were then used for RNA extraction and qPCR analysis to assess gene knockout efficiency.

### Bulk RNA sequencing and data analysis

HUVECs were infected with Ad-Moesin^WT^ or Ad-Moesin^T558D^ for 48 hrs. Total RNA was then isolated for RNA sequencing. RNA quality was assessed using an Agilent 2100 Bioanalyzer (Agilent Technologies). Sequencing was performed by Majorbio Bio-Pharm Technology Co. Ltd. (Shanghai, China). Differential gene expression between groups was analyzed using the DESeq2 package. The RNA-seq data generated in this study have been deposited in the Gene Expression Omnibus (GEO) database under accession code GSE302986.

### Membrane tension measurements by MSS

The membrane-bound tension biosensor MSS was kindly provided by Dr. Bo Liu (*40*). Specifically, MSS was constructed by PCR using a forward primer encoding 21 amino acids from Lyn kinase and a reverse primer encoding 14 amino acids from K-Ras, which anchor the tension sensor to phospholipids in lipid raft and non-lipid raft regions, respectively. MSS plasmids were transfected into EA.hy926 cells using Lipofectamine 3000 reagent (Invitrogen) according to the manufacturer’s instructions. FRET ratio images were generated by calculating YPet/ECFP using Ilastik and ImageJ.

### The oxygen-induced retinopathy (OIR) model

The OIR model was performed as previously described (*49*). Neonatal C57BL/6J mouse pups, together with their nursing mothers, were exposed to hyperoxia (75% O₂) from P7 to P12. Mice were then returned to room air. Tamoxifen (50 μL of 1 mg/mL) was administered daily from P12 to P14. On P17, mice were sacrificed and the retinas were harvested for analysis. Four radial incisions were made with spring scissors to flatten the whole-mount retinas, which were then imaged using a confocal microscope (LSM800, Carl Zeiss). Retinal neovascularization areas were quantified using ImageJ.

### Electroretinography (ERG)

ERG recordings were performed using the Celeris D430 rodent ERG testing system (Diagnosys LLC, MA, USA) as previously described (*68*). Briefly, mice from each group were dark-adapted for at least 8 hrs in a light-proof room prior to recording. Mice were then anesthetized, and pupils were dilated with one drop of 0.5% tropicamide (SINQI, China) for 5 mins.

Dark adaptation was maintained throughout the recording. The Celeris apparatus uses safe red (670 nm) and infrared (940 nm) LEDs for illumination. Mice were placed on a heated platform to maintain body temperature, and corneas were kept moist with 1–2% hydroxypropyl methylcellulose (Allergan, US). Two corneal electrodes with integrated stimulators were placed on the surfaces of the lubricated corneas. Impedances of the ground and reference electrodes were maintained between 1.5 and 2 kΩ during recording, and the impedances of the two corneal electrodes were kept between 2 and 5 kΩ. A stimulus intensity range of 0.01, 0.1, and 1 cd s/m² was used.

### Immunofluorescence of mouse tissues

For flat-mount retina immunofluorescence staining, eyeballs were harvested and fixed in 4% PFA/PBS for 1 hr at 4 °C. Retinas were dissected, permeabilized in PBS containing 1% Triton X-100 overnight at 4 °C, and then blocked with 0.5% Triton X-100 and 5% BSA for 3 hrs at room temperature. After blocking, retinas were incubated overnight at 4 °C with anti-Kindlin-2 antibody (Millipore, #MAB2617, 1:200), anti-p-ERM (CST, #3141S, 1:200), Ki67 (Abcam, #ab16667, 1:100), or ERG (Abcam, #ab92513, 1:100). After washing, retinas were incubated with Alexa Fluor 594-conjugated isolectin GS-IB4 (Invitrogen, #I21413, 1:100) and corresponding secondary antibodies for 3 hrs at room temperature.

For mouse embryo staining, embryos were harvested and immersion-fixed overnight in 4% PFA at 4 °C. After several washes with PBS, fixed embryos were transferred to 30% sucrose at 4 °C overnight, embedded in optimal cutting temperature (OCT) compound, and stored at –80 °C before sectioning with a cryostat (CM1950, Leica). Cryosections were washed, permeabilized in PBS containing 0.3% Triton X-100, blocked with 0.3% Triton X-100 and 3% BSA for 1 hr at room temperature, and incubated overnight at 4 °C with Alexa Fluor 594-conjugated isolectin GS-IB4 (Invitrogen, #I21413, 1:100). After washing, sections were incubated with the corresponding secondary antibody for 2 hrs at room temperature.

For immunofluorescence staining of mouse brain vibratome sections, brains were harvested and fixed in 4% PFA/PBS for 4 hrs at 4 °C. Samples were then embedded in 3% low-melting-point agarose in PBS for 30 mins at room temperature and sectioned using a vibratome (Leica, Germany). Vibratome sections were washed, permeabilized in PBS containing 0.3% Triton X-100, blocked in PBS containing 2% BSA and 0.3% Triton X-100 for 1 hr, and incubated overnight at 4 °C with Ter119 (R&D Systems, #MAB1125, 1:200). After washing, sections were incubated with Alexa Fluor 594-conjugated isolectin GS-IB4 (Invitrogen, #I21413, 1:100) or corresponding secondary antibodies for 2 hrs at room temperature. Images were acquired using a confocal microscope (LSM 800, Carl Zeiss).

### Immunofluorescence of cells in culture

Cells on glass coverslips were fixed with 2% PFA and permeabilized with 0.1% Triton X-100 in PBS for 30 mins. Samples were then blocked in PBS solution with 2%BSA, 0.1% TritonX-100 for 1 hr and stained with the primary antibody VEGFR2 (Cell SignalingTechennology, #2479S, 1:200) and secondary antibody. Images were acquired by confocal microscope (Zeiss LSM 800). More than 12 fields of view from at least three independent experiments were randomly chosen.

Cells grown on glass coverslips were fixed with 2% PFA and permeabilized with 0.1% Triton X-100 in PBS for 30 mins. Samples were then blocked in PBS containing 2% BSA and 0.1% Triton X-100 for 1 hr at room temperature, followed by staining with primary antibody VEGFR2 (Cell Signaling Technology, #2479S, 1:200) and the appropriate secondary antibody. Images were acquired using a confocal microscope (Zeiss LSM 800). For analysis, more than 12 random fields of view from at least three independent experiments were analyzed.

### Biotinylation assay

The biotinylation assay was carried out as previously described (*46*). HUVECs were grown to confluence and serum-starved for 8 hrs in M199 medium. Cells were rinsed and incubated with EZ-Link™ Sulfo-NHS-SS-Biotin (0.25 mg/mL; Thermo Scientific) in PBS at 4 °C for 1 hr. The reaction was quenched by washing the cells with 50 mM glycine in PBS. A portion of the cells was collected to assess total biotinylated surface proteins. The remaining cells were rinsed once with cold medium containing 1% BSA and then stimulated with VEGF (50 ng/mL) in M199 at 37 °C for various time points. After stimulation, cells were rinsed and incubated twice for 20 mins each on ice with the membrane-impermeable reducing agent glutathione (GSH, 45 mM; Sigma) in 75 mM NaCl, 75 mM NaOH, 1 mM EDTA, and 1% BSA. GSH was neutralized by two successive 5-minute incubations with iodoacetamide (5 mg/mL) in PBS. Cell lysates were prepared using NP-40 lysis buffer (Solarbio). Lysates were incubated overnight at 4 °C with 50 μL Pierce™ Streptavidin Magnetic Beads (Thermo, #88816). After washing, beads were resuspended in 2 × loading buffer, and samples were analyzed by western blotting using an anti-VEGFR2 antibody (CST, #2479S, 1:1000). The remaining lysates after bead incubation were used for immunoblotting of VEGFR2 (CST, #2479S, 1:1000), Kindlin-2 (Millipore, #MAB2617, 1:1000), and GAPDH (Utibody, #UM4002, 1:1000).

### Split GFP assay

Full-length Moesin was synthesized and cloned into the GFP1-10 plasmid, and full-length Kindlin-2 was synthesized and cloned into the GFP11 plasmid. The Moesin-GFP1-10 and Kindlin-2-GFP11 plasmids were then co-transfected into EA. Hy926 cells using Lipofectamine 3000 reagent (Invitrogen) according to the manufacturer’s protocol. Live-cell images were acquired using a confocal microscope (LSM 800, Carl Zeiss).

### BrdU incorporation assay in cells

The BrdU incorporation assay was performed as previously described (*69*). Briefly, HUVECs were cultured on collagen-coated coverslips in 24-well plates. After 48 hrs of siRNA transfection, BrdU (10 µM) was added and the cultures were incubated for 4 hrs at 37 °C. Cells were then fixed in 2% PFA/PBS for 20 mins and blocked in PBS containing 2% BSA and 0.3% Triton X-100 for 30 mins at room temperature. DNA denaturation was performed by incubating cells in ice-cold 0.1 M HCl for 20 mins, followed by 2 M HCl for 30 mins. Neutralization was carried out with sodium borate buffer (0.1 M Na_2_B_4_O_7_, pH 8.5) for 15 mins before primary antibody incubation. Cells were incubated overnight at 4 °C with an anti-BrdU antibody (Abcam, #ab6326, 1:100) in blocking solution, followed by incubation with the appropriate secondary antibody for 2 hrs at room temperature. Nuclei were counterstained with DAPI. Images were acquired using a confocal microscope (LSM 800, Carl Zeiss). Quantification was performed blind to the experimental condition.

### In vitro aortic ring assay

The aortic ring assay was performed as previously described (*49*). Briefly, 1-mm long segments of the thoracic aorta were cultured in Opti-MEM™ (Thermo Scientific, #31985070) supplemented with 100 U/ml penicillin and 100 µg/mL streptomycin overnight. The aortic rings were then embedded in growth factor-reduced Matrigel (Corning, #354230) in 24-well plates. Subsequently, the rings were cultured in Opti-MEM™ supplemented with 2.5% FBS in a 37 °C, 5% CO_2_ incubator for 5 days. After fixation in 4% PFA/PBS for 20 mins at room temperature, the aortic rings were permeabilized in PBS containing 0.3% Triton X-100, then blocked in PBS with 2% BSA and 0.3% Triton X-100 for 1 hr. Samples were incubated with Alexa Fluor 488-conjugated phalloidin (Sigma-Aldrich, 1:100) overnight at 4 °C. Images were acquired using a confocal microscope (LSM 800, Carl Zeiss).

### Fibrin gel bead sprouting assay

Cytodex 3 microcarrier beads (GE Healthcare) were pre-coated with HUVECs (including siRNA-transfected or adenovirus-infected variants) at a density of 200 cells per bead and subsequently embedded in fibrin gels (2 mg/mL fibrinogen in PBS, Calbiochem) polymerized with 0.625 U/mL thrombin (Sigma-Aldrich) and 0.15 U/mL aprotinin (Sigma-Aldrich). The embedded HUVEC-coated beads were cultured in M199 medium for 24 hrs, then fixed with 4% PFA for 15 mins, permeabilized with 0.2% Triton X-100, and blocked with 1% BSA in PBS. F-actin was stained using Alexa Fluor 488-conjugated phalloidin (Sigma-Aldrich, 1:100) for 2 hrs at room temperature. Images were acquired using a Zeiss LSM 800 confocal microscope, and quantitative analysis of sprout length per bead was performed using NIH ImageJ software.

### Recombinant adenovirus construction and infection

Recombinant adenoviruses expressing the wild-type *FERMT2* gene (GenBank accession no. NM_006832.3) and the wild-type *MSN* gene (GenBank accession no. NM_002444.3) were purchased from Shanghai Genechem. The Ad-Moesin^N62A^ adenovirus was generated by replacing the Asparagine at position 62 with Alanine and was produced by Shanghai Genechem. HUVECs were infected with adenovirus at a multiplicity of infection (MOI) of approximately 100. 48 hrs after infection, cells were harvested for further analysis.

### Recombinant AAV vector production and administration

Brain microvasculature endothelial cell–specific AAV-BR1 was generated as previously described (*50*). Briefly, HEK293T cells were transfected with transfer plasmids (pAAV-U6-MCS-shCtrl-zsGreen, pAAV-U6-MCS-sh*M*sn-zsGreen) along with pCapsid NRGTEWD (BR1) and pHelper. Transfection was performed using linear polyethylenimine (Polysciences). After five days, cells were harvested and lysed using salt-active nuclease. The virus was purified by iodixanol density-gradient ultracentrifugation, and the vector copy number was determined by qRT-PCR.

In the OIR model, recombinant AAV vectors were administrated via retro-orbital injection at a dose of 5×10 ^10^ gp per pup at P3.

### Statistical analysis

Statistical analyses were performed using GraphPad Prism version 9.0. All data are expressed as the mean ± SEM. To calculate statistical significance, a two-tailed Student’s t-test, one-way analysis of variance (ANOVA) followed by Tukey’s multiple comparisons were used. Sample size was determined according to previous publication where at least three animals per group were analyzed. The experimenters were blinded to animal genotype and grouping information and all data were derived from biological replicates as indicated. P-values less than 0.05 were considered statistically significant.

## Acknowledgments

The authors thank Marcus Fruttiger at UCL Institute of Ophthalmology (London, UK) for providing us the *Pdgfb-CreERT2* mice, Prof. Ju Chen at University of California, San Diego for providing the *Kindlin-2^fl/fl^* mice, Prof. Martin Trepel and Dr. Jakob Körbelin for providing the AAV-BR1 system. The authors thank Prof. Bo Liu at Dalian University of Technology for providing the MSS FRET probe. The authors thank the Core Facility of Research Center of Basic Medical Sciences at Tianjin Medical University for technical support.

## Author contributions

Conceptualization: X.W.

Funding Acquisition: J.Z. and X.W.

Investigation: L.W., Y.F., Z.Y., Y.L., T.Y., J.L., N.M. and Y.L

Methodology: L.W.

Supervision: K.O., K.Z, J.H, X.F., Y.S., J.Z. and X.W.

Writing-Original Draft: L.W. and X.W.

Writing-Review & Editing: All authors reviewed the paper and approved the final draft.

## Data and materials availability

All data needed to evaluate the conclusions in the paper are present in the paper and/or the Supplementary Materials. Bulk RNA sequencing data and single-cell RNA sequencing data are available in National Center for Biotechnology Information’s Gene Expression Omnibus under accession number GSE175895 (scRNA-seq on mouse retina from P6 and P10), GSE216676 (scRNA-seq on OIR model mouse retina from P17), GSE79306 (scRNA-seq on mouse brain from E11.5, E12.5, E13.5, E15.5, E16.5 and E17.5), GSE302986 (bulk RNA-seq on HUVEC infected with Ad-Moesin^WT^ and Ad-Moesin^T558D^), GSE 291008 (bulk RNA-seq on HRECs), GSE287078 (bulk RNA-seq on HUVECs), GSE281250 (bulk RNA-seq on HCMECs) and GSE94862 (bulk RNA-seq on purified P5 mouse brain ECs).

## References

1. H. G. Augustin, G. Y. Koh, Organotypic vasculature: From descriptive heterogeneity to functional pathophysiology. Science 357, eaal2379 (2017).

2. R. S. Apte, D. S. Chen, N. Ferrara, VEGF in Signaling and Disease: Beyond Discovery and Development. Cell 176, 1248–1264 (2019).

3. M. J. Davis, S. Earley, Y.-S. Li, S. Chien, Vascular mechanotransduction. Physiological Reviews 103, 1247–1421 (2023).

4. P. A. Galie, D. H. Nguyen, C. K. Choi, D. M. Cohen, P. A. Janmey, C. S. Chen, Fluid shear stress threshold regulates angiogenic sprouting. Proc Natl Acad Sci U S A 111, 7968–7973 (2014).

5. V. Gebala, R. Collins, I. Geudens, L. K. Phng, H. Gerhardt, Blood flow drives lumen formation by inverse membrane blebbing during angiogenesis in vivo. Nat Cell Biol 18, 443–450 (2016).

6. Y. Shen, X. Wang, J. Lu, M. Salfenmoser, N. M. Wirsik, N. Schleussner, A. Imle, A. Freire Valls, P. Radhakrishnan, J. Liang, G. Wang, T. Muley, M. Schneider, C. Ruiz de Almodovar, A. Diz-Munoz, T. Schmidt, Reduction of Liver Metastasis Stiffness Improves Response to Bevacizumab in Metastatic Colorectal Cancer. Cancer Cell 37, 800–817 e807 (2020).

7. F. Bordeleau, B. N. Mason, E. M. Lollis, M. Mazzola, M. R. Zanotelli, S. Somasegar, J. P. Califano, C. Montague, D. J. LaValley, J. Huynh, N. Mencia-Trinchant, Y. L. Negron Abril, D. C. Hassane, L. J. Bonassar, J. T. Butcher, R. S. Weiss, C. A. Reinhart-King, Matrix stiffening promotes a tumor vasculature phenotype. Proc Natl Acad Sci U S A 114, 492–497 (2017).

8. Y. Han, J. Yan, Z.-Y. Li, Y.-J. Fan, Z.-L. Jiang, J. Y. J. Shyy, S. Chien, Cyclic stretch promotes vascular homing of endothelial progenitor cells via Acsl1 regulation of mitochondrial fatty acid oxidation. Proceedings of the National Academy of Sciences 120, e2219630120 (2023).

9. M. Sotoudeh, Y.-S. Li, N. Yajima, C.-C. Chang, T.-C. Tsou, Y. Wang, S. Usami, A. Ratcliffe, S. Chien, J. Y. J. Shyy, Induction of apoptosis in vascular smooth muscle cells by mechanical stretch. American Journal of Physiology-Heart and Circulatory Physiology 282, H1709–H1716 (2002).

10. E. Rognoni, R. Ruppert, R. Fassler, The kindlin family: functions, signaling properties and implications for human disease. J Cell Sci 129, 17–27 (2016).

11. J. J. Dowling, E. Gibbs, M. Russell, D. Goldman, J. Minarcik, J. A. Golden, E. L. Feldman, Kindlin-2 is an essential component of intercalated discs and is required for vertebrate cardiac structure and function. Circ Res 102, 423–431 (2008).

12. E. Montanez, S. Ussar, M. Schifferer, M. Bosl, R. Zent, M. Moser, R. Fassler, Kindlin-2 controls bidirectional signaling of integrins. Genes Dev 22, 1325–1330 (2008).

13. L. Guo, C. Cui, K. Zhang, J. Wang, Y. Wang, Y. Lu, K. Chen, J. Yuan, G. Xiao, B. Tang, Y. Sun, C. Wu, Kindlin-2 links mechano-environment to proline synthesis and tumor growth. Nature Communications 10, 845 (2019).

14. X. Wu, Y. Lai, S. Chen, C. Zhou, C. Tao, X. Fu, J. Li, W. Tong, H. Tian, Z. Shao, C. Liu, D. Chen, X. Bai, H. Cao, G. Xiao, Kindlin-2 preserves integrity of the articular cartilage to protect against osteoarthritis. Nat Aging 2, 332–347 (2022).

15. H. Gao, L. Zhou, Y. Zhong, Z. Ding, S. Lin, X. Hou, X. Zhou, J. Shao, F. Yang, X. Zou, H. Cao, G. Xiao, Kindlin-2 haploinsufficiency protects against fatty liver by targeting Foxo1 in mice. Nature Communications 13, 1025 (2022).

16. X. Fu, B. Zhou, Q. Yan, C. Tao, L. Qin, X. Wu, S. Lin, S. Chen, Y. Lai, X. Zou, Z. Shao, M. Wang, D. Chen, W. Jin, Y. Song, H. Cao, G. Zhang, G. Xiao, Kindlin-2 regulates skeletal homeostasis by modulating PTH1R in mice. Signal Transduct Target Ther 5, 297 (2020).

17. A. Chronopoulos, S. D. Thorpe, E. Cortes, D. Lachowski, A. J. Rice, V. V. Mykuliak, T. Rog, D. A. Lee, V. P. Hytonen, A. E. Del Rio Hernandez, Syndecan-4 tunes cell mechanics by activating the kindlin-integrin-RhoA pathway. Nat Mater 19, 669–678 (2020).

18. N. Ma, F. Wu, J. Liu, Z. Wu, L. Wang, B. Li, Y. Liu, X. Dong, J. Hu, X. Fang, H. Zhang, D. Ai, J. Zhou, X. Wang, Kindlin-2 Phase Separation in Response to Flow Controls Vascular Stability. Circ Res 135, 1141–1160 (2024).

19. Y. Dong, G. Ma, X. Hou, Y. Han, Z. Ding, W. Tang, L. Chen, Y. Chen, B. Zhou, F. Rao, K. Lv, C. Du, H. Cao, Kindlin-2 controls angiogenesis through modulating Notch1 signaling. Cell Mol Life Sci 80, 223 (2023).

20. E. Sitarska, A. Diz-Munoz, Pay attention to membrane tension: Mechanobiology of the cell surface. Current Opinion in Cell Biology 66, 11–18 (2020).

21. D. Raucher, T. Stauffer, W. Chen, K. Shen, S. Guo, J. D. York, M. P. Sheetz, T. Meyer, Phosphatidylinositol 4,5-Bisphosphate Functions as a Second Messenger that Regulates Cytoskeleton–Plasma Membrane Adhesion. Cell 100, 221–228 (2000).

22. B. Rouven Brückner, A. Pietuch, S. Nehls, J. Rother, A. Janshoff, Ezrin is a Major Regulator of Membrane Tension in Epithelial Cells. Scientific Reports 5, 14700 (2015).

23. W. A. Harris, A. Diz-Muñoz, M. Krieg, M. Bergert, I. Ibarlucea-Benitez, D. J. Muller, E. Paluch, C.-P. Heisenberg, Control of Directed Cell Migration In Vivo by Membrane-to-Cortex Attachment. PLoS Biology 8, e1000544 (2010).

24. M. Berryman, Z. Franck, A. Bretscher, Ezrin is concentrated in the apical microvilli of a wide variety of epithelial cells whereas moesin is found primarily in endothelial cells. Journal of Cell Science 105, 1025–1043 (1993).

25. P. Vitorino, S. Yeung, A. Crow, J. Bakke, T. Smyczek, K. West, E. McNamara, J. Eastham-Anderson, S. Gould, S. F. Harris, C. Ndubaku, W. Ye, MAP4K4 regulates integrin-FERM binding to control endothelial cell motility. Nature 519, 425–430 (2015).

26. Y. Wang, M. S. Kaiser, J. D. Larson, A. Nasevicius, K. J. Clark, S. A. Wadman, S. E. Roberg-Perez, S. C. Ekker, P. B. Hackett, M. McGrail, J. J. Essner, Moesin1 and Ve-cadherin are required in endothelial cells during in vivo tubulogenesis. Development 137, 3119–3128 (2010).

27. C. Wu, H. Jiao, Y. Lai, W. Zheng, K. Chen, H. Qu, W. Deng, P. Song, K. Zhu, H. Cao, D. L. Galson, J. Fan, H.-J. Im, Y. Liu, J. Chen, D. Chen, G. Xiao, Kindlin-2 controls TGF-β signalling and Sox9 expression to regulate chondrogenesis. Nature Communications 6, ncomms8531 (2015).

28. S. Claxton, V. Kostourou, S. Jadeja, P. Chambon, K. Hodivala-Dilke, M. Fruttiger, Efficient, inducible Cre-recombinase activation in vascular endothelium. genesis 46, 74–80 (2008).

29. X. Wang, A. Freire Valls, G. Schermann, Y. Shen, I. M. Moya, L. Castro, S. Urban, G. M. Solecki, F. Winkler, L. Riedemann, R. K. Jain, M. Mazzone, T. Schmidt, T. Fischer, G. Halder, C. Ruiz de Almodóvar, YAP/TAZ Orchestrate VEGF Signaling during Developmental Angiogenesis. Developmental Cell 42, 462–478 (2017).

30. M. Fruttiger, Development of the retinal vasculature. Angiogenesis 10, 77–88 (2007).

31. J. H. Bae, M. J. Yang, S.-h. Jeong, J. Kim, S. P. Hong, J. W. Kim, Y. H. Kim, G. Y. Koh, Gatekeeping role of Nf2/Merlin in vascular tip EC induction through suppression of VEGFR2 internalization. Science Advances 8, abn2611 (2022).

32. H. Yamamoto, M. Ehling, K. Kato, K. Kanai, M. van Lessen, M. Frye, D. Zeuschner, M. Nakayama, D. Vestweber, R. H. Adams, Integrin β1 controls VE-cadherin localization and blood vessel stability. Nature Communications 6, ncomms7429 (2015).

33. M. G. Romei, S. G. Boxer, Split Green Fluorescent Proteins: Scope, Limitations, and Outlook. Annual Review of Biophysics 48, 19–44 (2019).

34. I. Paredes, P. Himmels, C. Ruiz de Almodóvar, Neurovascular Communication during CNS Development. Developmental Cell 45, 10–32 (2018).

35. M. A. Pearson, D. Reczek, A. Bretscher, P. A. Karplus, Structure of the ERM Protein Moesin Reveals the FERM Domain Fold Masked by an Extended Actin Binding Tail Domain. Cell 101, 259–270 (2000).

36. B. T. Fievet, A. Gautreau, C. Roy, L. Del Maestro, P. Mangeat, D. Louvard, M. Arpin, Phosphoinositide binding and phosphorylation act sequentially in the activation mechanism of ezrin. The Journal of Cell Biology 164, 653–659 (2004).

37. M. Baumgartner, A. L. Sillman, E. M. Blackwood, J. Srivastava, N. Madson, J. W. Schilling, J. H. Wright, D. L. Barber, The Nck-interacting kinase phosphorylates ERM proteins for formation of lamellipodium by growth factors. Proceedings of the National Academy of Sciences 103, 13391–13396 (2006).

38. K. Ben-Aissa, G. Patino-Lopez, N. V. Belkina, O. Maniti, T. Rosales, J.-J. Hao, M. J. Kruhlak, J. R. Knutson, C. Picart, S. Shaw, Activation of Moesin, a Protein That Links Actin Cytoskeleton to the Plasma Membrane, Occurs by Phosphatidylinositol 4,5-bisphosphate (PIP2) Binding Sequentially to Two Sites and Releasing an Autoinhibitory Linker. Journal of Biological Chemistry 287, 16311–16323 (2012).

39. A. Bretscher, K. Edwards, R. G. Fehon, ERM proteins and merlin: integrators at the cell cortex. Nature Reviews Molecular Cell Biology 3, 586–599 (2002).

40. W. Li, X. Yu, F. Xie, B. Zhang, S. Shao, C. Geng, A. u. R. Aziz, X. Liao, B. Liu, A Membrane-Bound Biosensor Visualizes Shear Stress-Induced Inhomogeneous Alteration of Cell Membrane Tension. iScience 7, 180–190 (2018).

41. S. Wang, b. Bian, W. Shi, T. M€oller, R. I. Stegmeyer, B. Strilic, T. Li, Z. Yuan, C. Wang, N. Wettschureck, D. Vestweber, S. Offermanns, Mechanosensation by endothelial PIEZO1 is required for leukocyte diapedesis. BLOOD 140, 171–183 (2022).

42. M. Simons, E. Gordon, L. Claesson-Welsh, Mechanisms and regulation of endothelial VEGF receptor signalling. Nature Reviews Molecular Cell Biology 17, 611–625 (2016).

43. J. J. Thottacherry, A. J. Kosmalska, A. Kumar, A. S. Vishen, A. Elosegui-Artola, S. Pradhan, S. Sharma, P. P. Singh, M. C. Guadamillas, N. Chaudhary, R. Vishwakarma, X. Trepat, M. A. del Pozo, R. G. Parton, M. Rao, P. Pullarkat, P. Roca-Cusachs, S. Mayor, Mechanochemical feedback control of dynamin independent endocytosis modulates membrane tension in adherent cells. Nature Communications 9, 4217 (2018).

44. J. Dai, M. P. Sheetz, Regulation of Endocytosis, Exocytosis, and Shape by Membrane Tension. Cold Spring Harbor Symposia on Quantitative Biology 60, 567–571 (1995).

45. H. De Belly, A. Stubb, A. Yanagida, C. Labouesse, P. H. Jones, E. K. Paluch, K. J. Chalut, Membrane Tension Gates ERK-Mediated Regulation of Pluripotent Cell Fate. Cell Stem Cell 28, 273–284 (2021).

46. G. Genet, K. Boyé, T. Mathivet, R. Ola, F. Zhang, A. Dubrac, J. Li, N. Genet, L. Henrique Geraldo, L. Benedetti, S. Künzel, L. Pibouin-Fragner, J.-L. Thomas, A. Eichmann, Endophilin-A2 dependent VEGFR2 endocytosis promotes sprouting angiogenesis. Nature Communications 10, 2350 (2019).

47. M. Nakayama, A. Nakayama, M. van Lessen, H. Yamamoto, S. Hoffmann, H. C. A. Drexler, N. Itoh, T. Hirose, G. Breier, D. Vestweber, J. A. Cooper, S. Ohno, K. Kaibuchi, R. H. Adams, Spatial regulation of VEGF receptor endocytosis in angiogenesis. Nature Cell Biology 15, 249–260 (2013).

48. C. Roffay, G. Molinard, K. Kim, M. Urbanska, V. Andrade, V. Barbarasa, P. Nowak, V. Mercier, J. García-Calvo, S. Matile, R. Loewith, A. Echard, J. Guck, M. Lenz, A. Roux, Passive coupling of membrane tension and cell volume during active response of cells to osmosis. Proceedings of the National Academy of Sciences 118, e2103228118 (2021).

49. Y. Lei, Q. Liu, B. Chen, F. Wu, Y. Li, X. Dong, N. Ma, Z. Wu, Y. Zhu, L. Wang, Y. Fu, Y. Liu, Y. Song, M. Du, H. Zhang, J. Zhu, T. J. Lyons, T. Wang, J. Hu, H. Xu, M. Chen, H. Yan, X. Wang, Protein O-GlcNAcylation coupled to Hippo signaling drives vascular dysfunction in diabetic retinopathy. Nature Communications 15, 9334 (2024).

50. J. Körbelin, G. Dogbevia, S. Michelfelder, D. A. Ridder, A. Hunger, J. Wenzel, H. Seismann, M. Lampe, J. Bannach, M. Pasparakis, J. A. Kleinschmidt, M. Schwaninger, M. Trepel, A brain microvasculature endothelial cell-specific viral vector with the potential to treat neurovascular and neurological diseases. EMBO Molecular Medicine 8, 609–625 (2016).

51. X. Dong, Y. Song, Y. Liu, X. Kou, T. Yang, S. X. Shi, K. He, Y. Li, Z. Li, X. Yao, J. Guo, B. Cui, Z. Wu, Y. Lei, M. Du, M. Chen, H. Xu, Q. Liu, F.-D. Shi, X. Wang, H. Yan, Natural killer cells promote neutrophil extracellular traps and restrain macular degeneration in mice. Science Translational Medicine 16, eadi6626 (2024).

52. T. Mammoto, D. E. Ingber, Mechanical control of tissue and organ development. Development 137, 1407–1420 (2010).

53. P.-A. Pouille, P. Ahmadi, A.-C. Brunet, E. Farge, Mechanical Signals Trigger Myosin II Redistribution and Mesoderm Invagination in Drosophila Embryos. Science Signaling 2, 1–9 (2009).

54. J. M. Muncie, V. M. Weaver, Membrane Tension Locks In Pluripotency. Cell Stem Cell 28, 175–176 (2021).

55. M. G. Lampugnani, F. Orsenigo, M. C. Gagliani, C. Tacchetti, E. Dejana, Vascular endothelial cadherin controls VEGFR-2 internalization and signaling from intracellular compartments. The Journal of Cell Biology 174, 593–604 (2006).

56. E. Boucrot, A. P. A. Ferreira, L. Almeida-Souza, S. Debard, Y. Vallis, G. Howard, L. Bertot, N. Sauvonnet, H. T. McMahon, Endophilin marks and controls a clathrin-independent endocytic pathway. Nature 517, 460–465 (2014).

57. D. Basagiannis, S. Zografou, C. Murphy, T. Fotsis, L. Morbidelli, M. Ziche, C. Bleck, J. Mercer, S. Christoforidis, VEGF induces signalling and angiogenesis by directing VEGFR2 internalisation through macropinocytosis. Journal of Cell Science 129, 4091–4104 (2016).

58. T. Wu, B. Zhang, F. Ye, Z. Xiao, A potential role for caveolin-1 in VEGF-induced fibronectin upregulation in mesangial cells: involvement of VEGFR2 and Src. American Journal of Physiology-Renal Physiology 304, F820–F830 (2013).

59. H. J. Zhou, L. Qin, Q. Jiang, K. N. Murray, H. Zhang, B. Li, Q. Lin, M. Graham, X. Liu, J. Grutzendler, W. Min, Caveolae-mediated Tie2 signaling contributes to CCM pathogenesis in a brain endothelial cell-specific Pdcd10-deficient mouse model. Nature Communications 12, 504 (2021).

60. A. Salikhova, L. Wang, A. A. Lanahan, M. Liu, M. Simons, W. P. J. Leenders, D. Mukhopadhyay, A. Horowitz, Vascular Endothelial Growth Factor and Semaphorin Induce Neuropilin-1 Endocytosis via Separate Pathways. Circulation Research 103, e71–79 (2008).

61. J. H. R. Hetmanski, H. de Belly, I. Busnelli, T. Waring, R. V. Nair, V. Sokleva, O. Dobre, A. Cameron, N. Gauthier, C. Lamaze, J. Swift, A. del Campo, T. Starborg, T. Zech, J. G. Goetz, E. K. Paluch, J.-M. Schwartz, P. T. Caswell, Membrane Tension Orchestrates Rear Retraction in Matrix-Directed Cell Migration. Developmental Cell 51, 460–475 (2019).

62. M. Sixt, A. Diz-Muñoz, K. Thurley, S. Chintamen, S. J. Altschuler, L. F. Wu, D. A. Fletcher, O. D. Weiner, Membrane Tension Acts Through PLD2 and mTORC2 to Limit Actin Network Assembly During Neutrophil Migration. PLOS Biology 14, e1002474 (2016).

63. K. Tsujita, T. Takenawa, T. Itoh, Feedback regulation between plasma membrane tension and membrane-bending proteins organizes cell polarity during leading edge formation. Nature Cell Biology 17, 749–758 (2015).

64. J. Mueller, G. Szep, M. Nemethova, I. de Vries, A. D. Lieber, C. Winkler, K. Kruse, J. V. Small, C. Schmeiser, K. Keren, R. Hauschild, M. Sixt, Load Adaptation of Lamellipodial Actin Networks. Cell 171, 188–200 (2017).

65. Y. Wang, M. Nakayama, M. E. Pitulescu, T. S. Schmidt, M. L. Bochenek, A. Sakakibara, S. Adams, A. Davy, U. Deutsch, U. Lüthi, A. Barberis, L. E. Benjamin, T. Mäkinen, C. D. Nobes, R. H. Adams, Ephrin-B2 controls VEGF-induced angiogenesis and lymphangiogenesis. Nature 465, 483–486 (2010).

66. Y. Lei, J. Hu, J. Zhao, Q. Liu, S. W. Zhang, F. Wu, Y. Liu, H. Ren, X. Qin, X. Wu, F. Gao, J. Hu, K. Ouyang, Q. Liu, X. Zheng, L. Shi, X. Wang, Deubiquitinase USP9X controls Wnt signaling for CNS vascular formation and barrier maintenance. Developmental Cell, 1618-1635.e1617 (2025).

67. J. H. Bae, M. J. Yang, S.-h. Jeong, J. Kim, S. P. Hong, J. W. Kim, Y. H. Kim, G. Y. Koh, Gatekeeping role of Nf2/Merlin in vascular tip EC induction through suppression of VEGFR2 internalization. Science Advances 8, 1–19 (2022).

68. K. He, X. Dong, T. Yang, Z. Li, Y. Liu, J. He, M. Wu, S. Wei-Zhang, P. Kaysar, B. Cui, X. Yao, L. Zhang, W. Zhou, H. Xu, J. Wei, Q. Liu, J. Hu, X. Wang, H. Yan, Smoking aggravates neovascular age-related macular degeneration via Sema4D-PlexinB1 axis-mediated activation of pericytes. Nature Communications 16, 2821 (2025).

69. X. Jiang, J. Hu, Z. Wu, S. T. Cafarello, M. Di Matteo, Y. Shen, X. Dong, H. Adler, M. Mazzone, C. Ruiz de Almodovar, X. Wang, Protein Phosphatase 2A Mediates YAP Activation in Endothelial Cells Upon VEGF Stimulation and Matrix Stiffness. Frontiers in Cell and Developmental Biology 9, 675562 (2021).

