## Supplemental figure and table for "Kindlin-2-Moesin interaction orchestrates sprouting angiogenesis via modulating endothelial membrane mechanics and VEGF signaling"

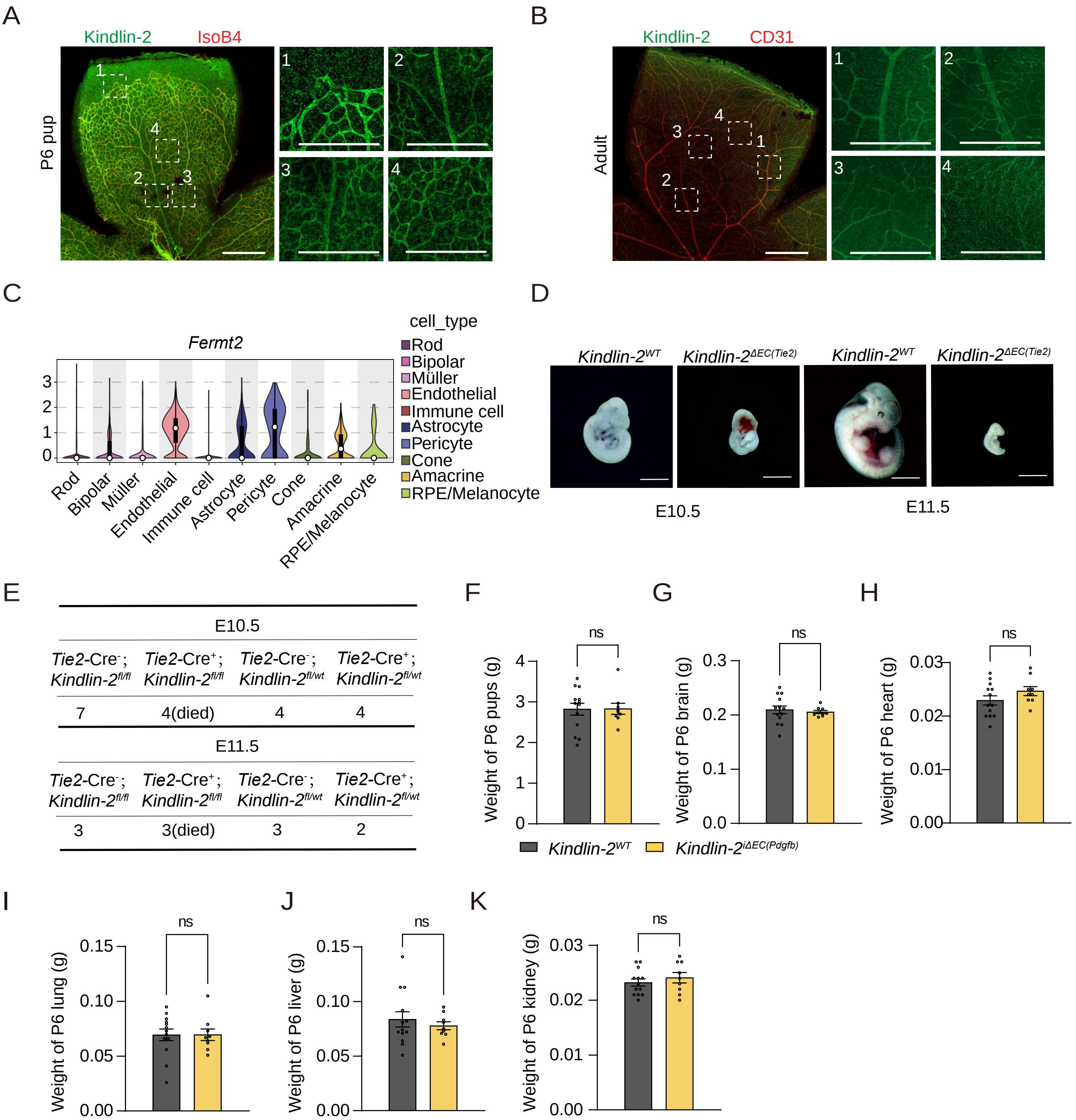


| E10.5 | | | |
|---|---|---|---|
| *Tie2*-Cre$^-$; *Kindlin-2*$^{fl/fl}$ | *Tie2*-Cre$^+$; *Kindlin-2*$^{fl/fl}$ | *Tie2*-Cre$^-$; *Kindlin-2*$^{fl/wt}$ | *Tie2*-Cre$^+$; *Kindlin-2*$^{fl/wt}$ |
| 7 | 4(died) | 4 | 4 |
| **E11.5** | | | |
| *Tie2*-Cre$^-$; *Kindlin-2*$^{fl/fl}$ | *Tie2*-Cre$^+$; *Kindlin-2*$^{fl/fl}$ | *Tie2*-Cre$^-$; *Kindlin-2*$^{fl/wt}$ | *Tie2*-Cre$^+$; *Kindlin-2*$^{fl/wt}$ |
| 3 | 3(died) | 3 | 2 |

Figure S1

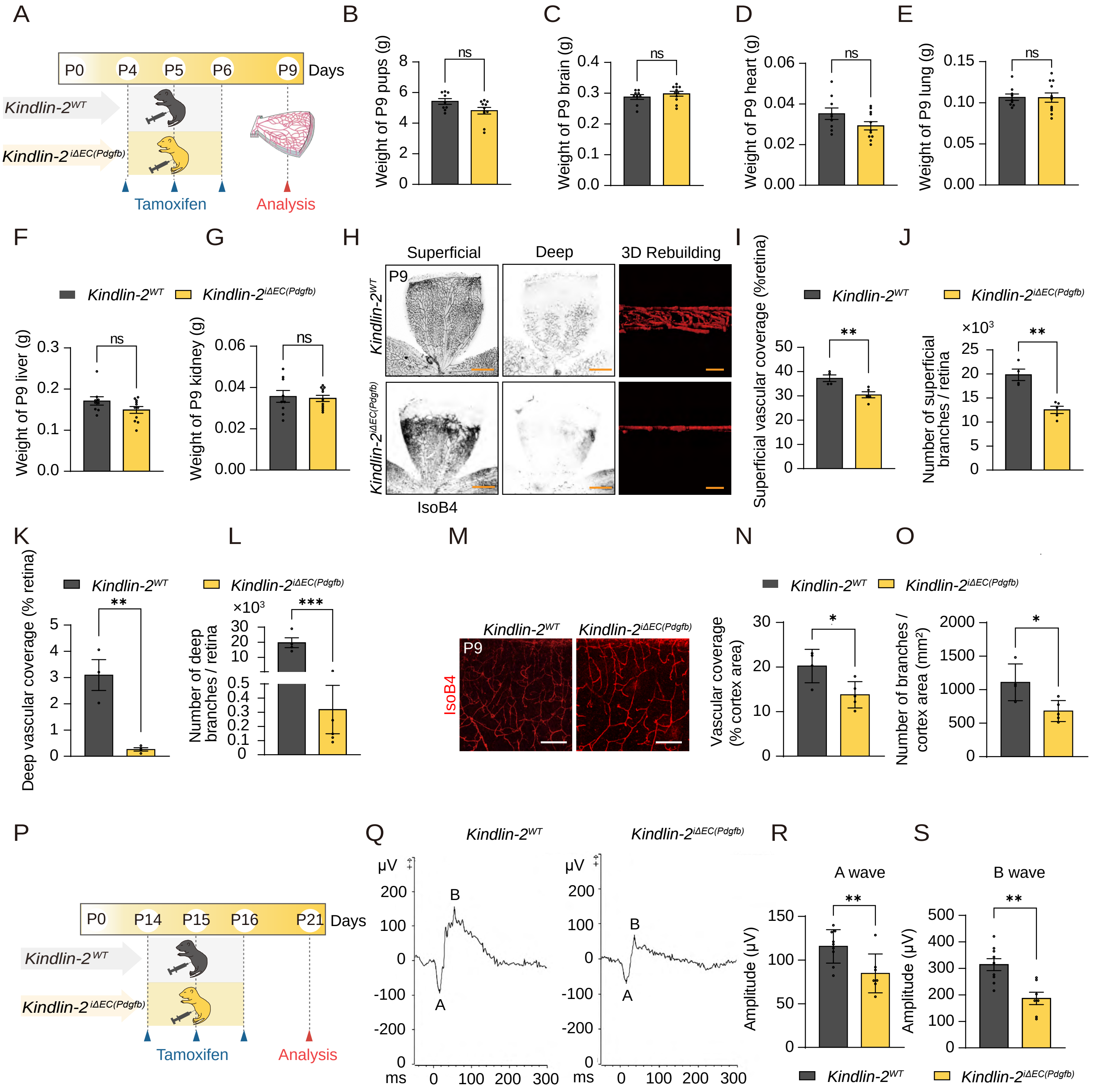

A
P0 P4 P5 P6 P9 Days
Kindlin-2WT
Kindlin-2iΔEC(Pdgfb)
Tamoxifen
Analysis
B
Weight of P9 pups (g)
ns
C
Weight of P9 brain (g)
ns
D
Weight of P9 heart (g)
ns
E
Weight of P9 lung (g)
ns
F
Weight of P9 liver (g)
ns
G
Weight of P9 kidney (g)
ns
H
Superficial
Deep
3D Rebuilding
P9
IsoB4
I
Superficial vascular coverage (%retina)
**
J
×10³
Number of superficial branches / retina
**
K
Deep vascular coverage (% retina)
**
L
×10³
Number of deep branches / retina
***
M
P9
IsoB4
N
Vascular coverage (% cortex area)
*
O
Number of branches / cortex area (mm²)
*
P
P0 P14 P15 P16 P21 Days
Tamoxifen
Analysis
Q
μV
ms
B
A
R
A wave
Amplitude (μV)
**
S
B wave
Amplitude (μV)
**


Figure S2

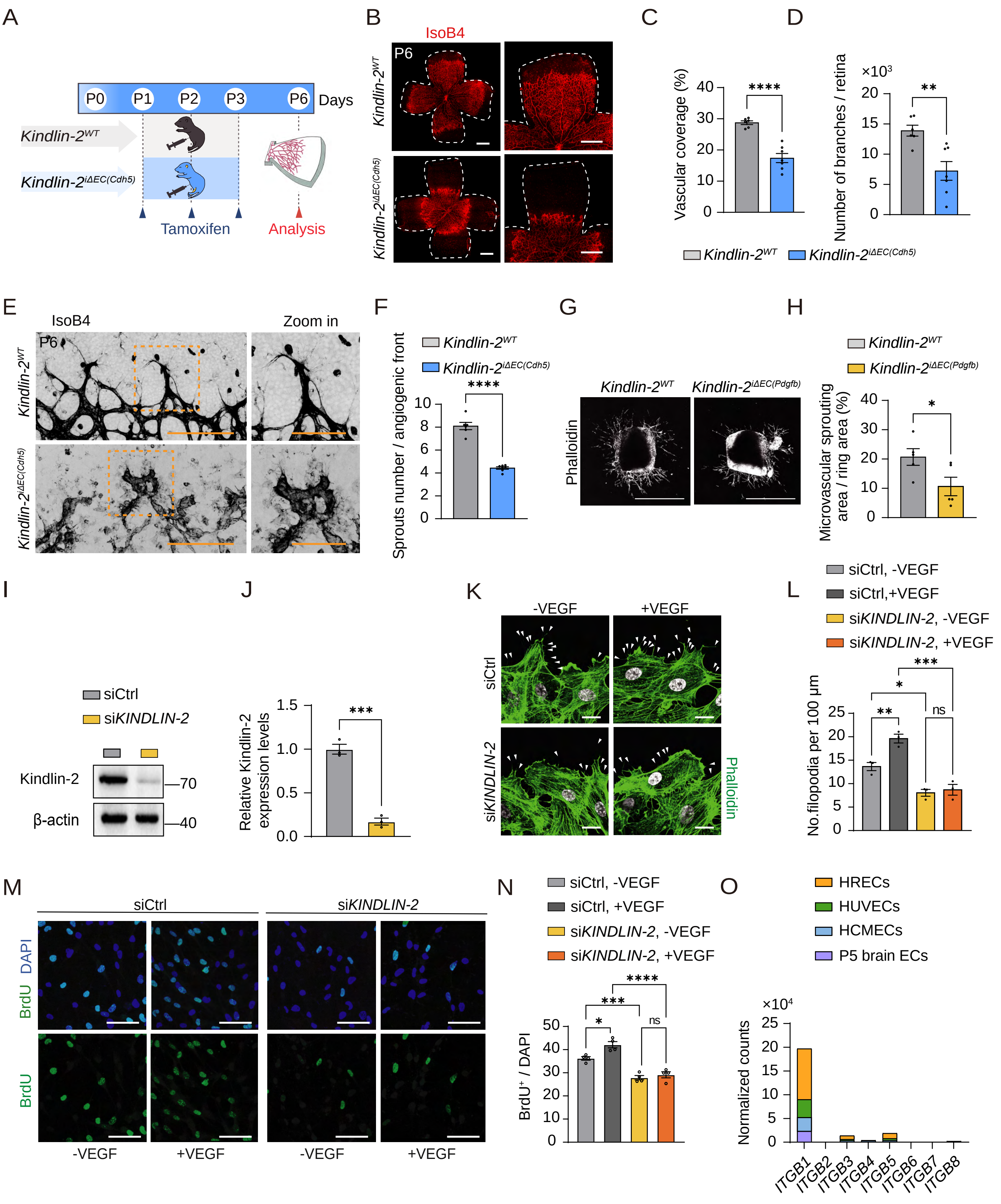

A
P0 P1 P2 P3 P6 Days
Kindlin-2WT
Kindlin-2iΔEC(Cdh5)
Tamoxifen
Analysis
B
IsoB4
P6
Kindlin-2WT
Kindlin-2iΔEC(Cdh5)
C
Vascular coverage (%)
D
Number of branches / retina
×10³
Kindlin-2WT
Kindlin-2iΔEC(Cdh5)
E
IsoB4
Zoom in
F
Sprouts number / angiogenic front
G
Phalloidin
Kindlin-2iΔEC(Pdgfb)
H
Microvascular sprouting area / ring area (%)
I
siCtrl
siKINDLIN-2
Kindlin-2
β-actin
70
40
J
Relative Kindlin-2 expression levels
K
-VEGF
+VEGF
L
siCtrl, -VEGF
siCtrl,+VEGF
siKINDLIN-2, -VEGF
siKINDLIN-2, +VEGF
No.filopodia per 100 µm
M
DAPI
BrdU
N
BrdU+ / DAPI
O
HRECs
HUVECs
HCMECs
P5 brain ECs
×10⁴
Normalized counts
ITGB1 ITGB2 ITGB3 ITGB4 ITGB5 ITGB6 ITGB7 ITGB8


Figure S3

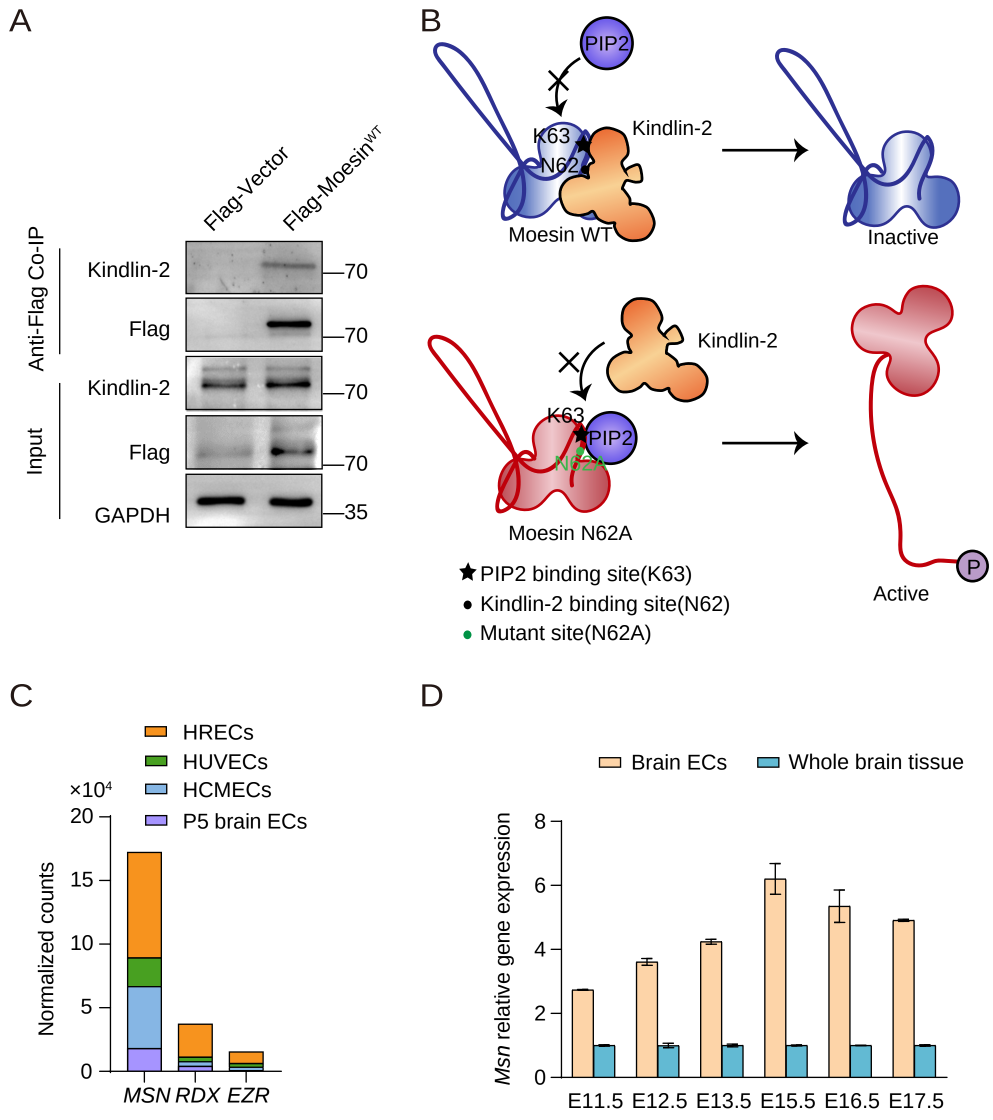

A
Flag-Vector
Flag-Moesin^WT
Anti-Flag Co-IP
Input
Kindlin-2
Flag
Kindlin-2
Flag
GAPDH
70
70
70
70
35
B
PIP2
K63
N62
Kindlin-2
Moesin WT
Inactive
Kindlin-2
K63
PIP2
N62A
Moesin N62A
P
Active
PIP2 binding site(K63)
Kindlin-2 binding site(N62)
Mutant site(N62A)
C
HRECs
HUVECs
HCMECs
P5 brain ECs
×10⁴
Normalized counts
20
15
10
5
0
MSN
RDX
EZR
D
Brain ECs
Whole brain tissue
Msn relative gene expression
8
6
4
2
0
E11.5
E12.5
E13.5
E15.5
E16.5
E17.5


Figure S4

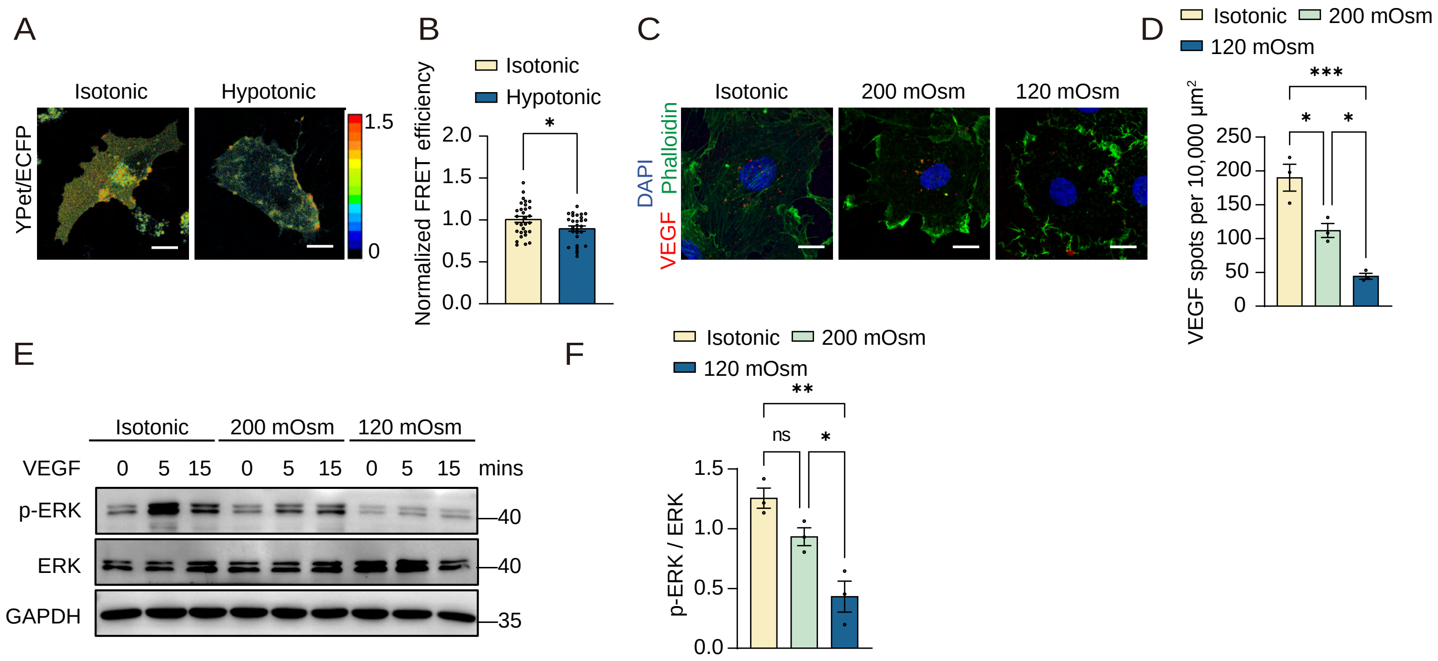

A
Isotonic
Hypotonic
YPet/ECFP
1.5
0
B
Isotonic
Hypotonic
Normalized FRET efficiency
2.0
1.5
1.0
0.5
0.0
*
C
Isotonic
200 mOsm
120 mOsm
DAPI
Phalloidin
VEGF
D
Isotonic
200 mOsm
120 mOsm
VEGF spots per 10,000 μm²
250
200
150
100
50
0
***
*
*
E
Isotonic
200 mOsm
120 mOsm
VEGF
0
5
15
0
5
15
0
5
15
mins
p-ERK
ERK
GAPDH
40
40
35
F
Isotonic
200 mOsm
120 mOsm
p-ERK / ERK
1.5
1.0
0.5
0.0
**
ns
*


Figure S5

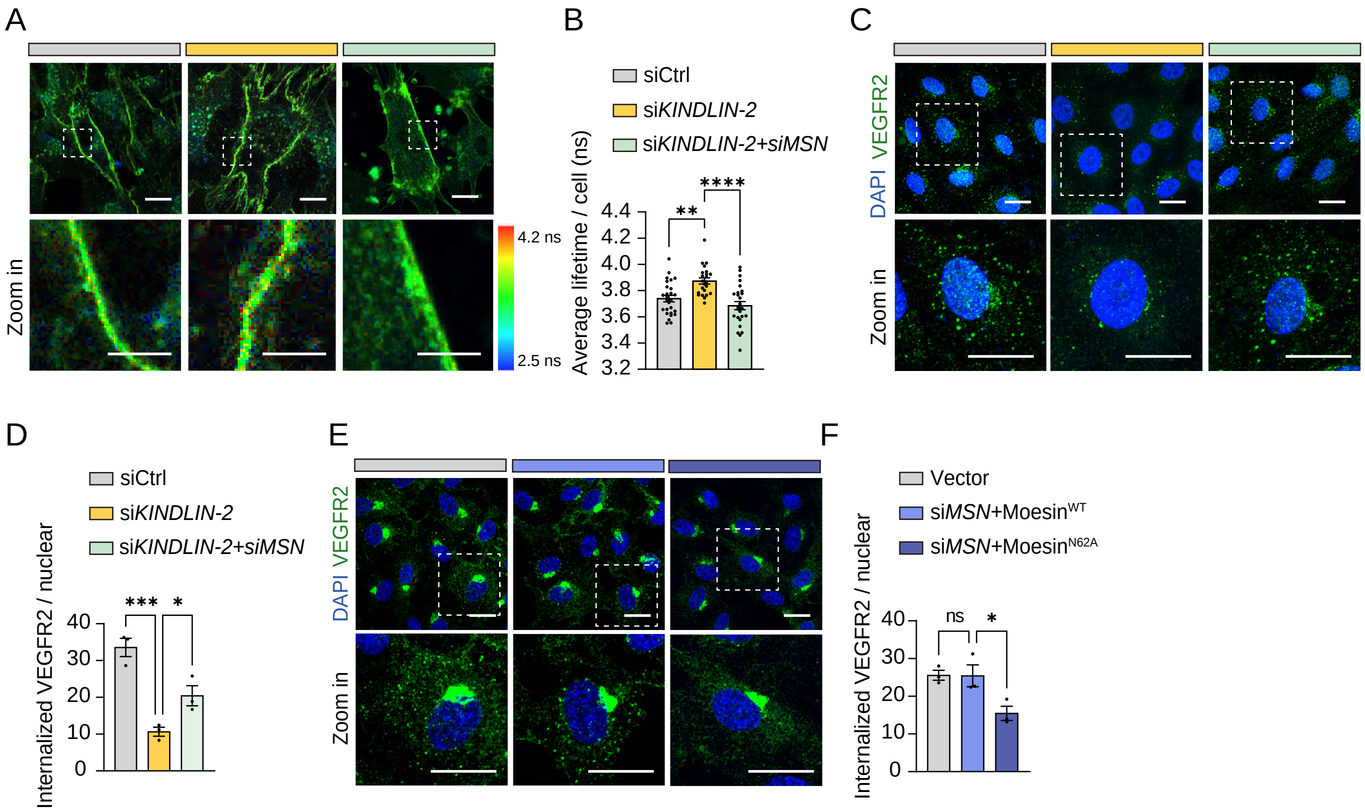

A
Zoom in
4.2 ns
2.5 ns
B
siCtrl
siKINDLIN-2
siKINDLIN-2+siMSN
Average lifetime / cell (ns)
4.4
4.2
4.0
3.8
3.6
3.4
3.2
**
****
C
DAPI VEGFR2
Zoom in
D
siCtrl
siKINDLIN-2
siKINDLIN-2+siMSN
Internalized VEGFR2 / nuclear
40
30
20
10
0
***
*
E
DAPI VEGFR2
Zoom in
F
Vector
siMSN+MoesinWT
siMSN+MoesinN62A
Internalized VEGFR2 / nuclear
40
30
20
10
0
ns
*


Figure S6

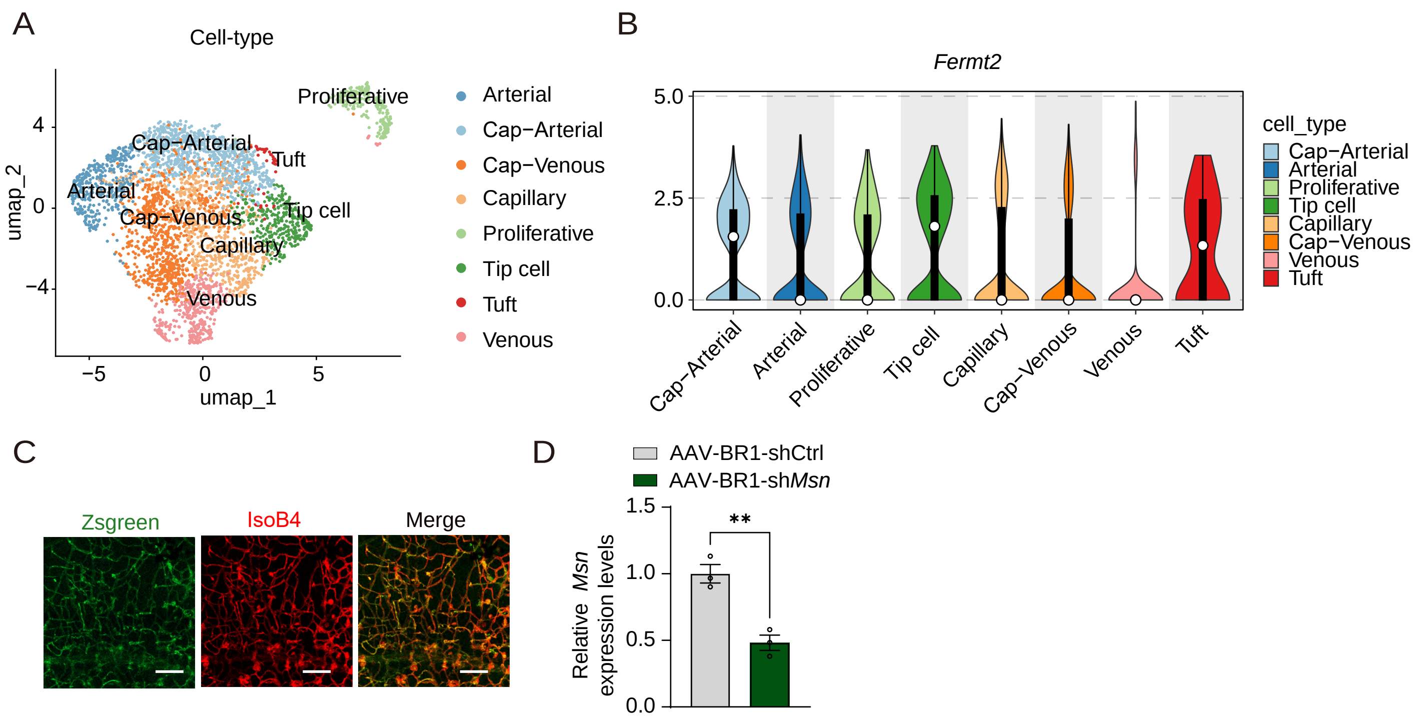
A
Cell-type
Proliferative
Cap−Arterial
Tuft
Arterial
Cap−Venous
Tip cell
Capillary
Venous
umap_2
umap_1
Arterial
Cap−Arterial
Cap−Venous
Capillary
Proliferative
Tip cell
Tuft
Venous
B
Fermt2
cell_type
Cap−Arterial
Arterial
Proliferative
Tip cell
Capillary
Cap−Venous
Venous
Tuft
C
Zsgreen
IsoB4
Merge
D
AAV-BR1-shCtrl
AAV-BR1-shMsn
Relative Msn expression levels
**

Figure S7

**Figure S1. Endothelial Kindlin-2 is essential for development**

(A) Flat-mounted immunofluorescence staining of Kindlin-2 (green) and IsoB4 (blood vessels, red) in postnatal day 6 (P6) pup retina. Kindlin-2 is highly expressed at the angiogenic front compared with the vascular plexus.

(B) Flat-mounted immunofluorescence staining of Kindlin-2 (green) and CD31 (red) in adult mouse retina.

(C) Violin plot showing the expression levels of Kindlin-2 (encoded by *Fermt2*) across different cell types in P10 mouse retina (data were analyzed from GSE175895).

(D) Representative images of *Kindlin-2*$^{WT}$ (*Tie2*-Cre$^{-}$; *Kindlin-2*$^{fl/fl}$) and *Kindlin-2*$^{\Delta EC(Tie2)}$ (*Tie2*-Cre$^{+}$; *Kindlin-2*$^{fl/fl}$) embryos at E10.5 and E11.5.

(E) Genotype distribution of embryos collected at E10.5 and E11.5 from the mating of *Tie2*-Cre$^{+}$; *Kindlin-2*$^{fl/wt}$ males with *Tie2*-Cre$^{-}$; *Kindlin-2*$^{fl/fl}$ females. Note that all embryos in *Tie2*-Cre$^{+}$; *Kindlin-2*$^{fl/fl}$ group were dead upon collection.

(D–I) Body weight of P6 *Kindlin-2*$^{WT}$ and *Kindlin-2*$^{i\Delta EC(Pdgfb)}$ pups, and weights of the brain, heart, lung, liver, and kidney (n = 13/9 pups).

Data are presented as mean ± SEM. ns represent no significant difference, two-tailed Student's t-test. Scale bars: 1 mm in (D), 500 μm in low magnification of (A) and (B), 200 μm in high magnification of (A) and (B).

**Figure S2. Endothelial Kindlin-2 is required for retinal and brain vascularization**

(A) Schematic illustration of the Tamoxifen administration schedule for Kindlin-$2^{WT}$ and *Kindlin-*$2^{i\Delta EC(Pdgfb)}$ mice. Tamoxifen was administered from P4 to P6, and retinas were analyzed at P9.

(B–G) Quantification of body weight and organ weights (brain, heart, liver, lung, and kidney) of P9 *Kindlin-*$2^{WT}$ and *Kindlin-*$2^{i\Delta EC(Pdgfb)}$ pups (n = 9/11 pups).

(H) Left: representative confocal images of the superficial vascular layer in P9 *Kindlin-*$2^{WT}$ and *Kindlin-*$2^{i\Delta EC(Pdgfb)}$ retinas. Middle: representative images of the deep vascular layer. Right: 3D reconstruction of retinal vasculature. Vessels stained with IsoB4 (red).

(I–L) Quantification of (H), showing reduced vascular coverage and fewer branch points in both superficial and deep vascular layers (n = 4/5 pups).

(M) Representative confocal images of brain sections from P9 *Kindlin-*$2^{WT}$ and *Kindlin-*$2^{i\Delta EC(Pdgfb)}$ pups stained with IsoB4 (red).

(N–O) Quantification of (M), showing reduced vascular coverage and branch points in *Kindlin-*$2^{i\Delta EC(Pdgfb)}$ brains (n = 4/5 pups).

(P) Schematic illustration of the Tamoxifen administration schedule for P21 *Kindlin-*$2^{WT}$ and *Kindlin-*$2^{i\Delta EC(Pdgfb)}$ mice. Tamoxifen was administered from P14 to P16.

(Q) Electroretinography of *Kindlin-*$2^{WT}$ and *Kindlin-*$2^{i\Delta EC(Pdgfb)}$ mice. Normal scotopic A- and B-wave responses were observed in *Kindlin-*$2^{WT}$ mice (left), whereas *Kindlin-*$2^{i\Delta EC(Pdgfb)}$ mice exhibited impaired responses (right).

(R–S) Quantification of scotopic A- and B-wave amplitudes in *Kindlin-*$2^{WT}$ and *Kindlin-*$2^{i\Delta EC(Pdgfb)}$ mice (n = 9/9 pups).

Data are presented as mean ± SEM. * $P < 0.05$; ** $P < 0.01$; *** $P < 0.001$, two-tailed Student's t-test. Scale bars: 50 μm in right panel of (H); 100 μm in (M); 500 μm in left and middle panel of (H).

**Figure S3. Loss of Kindlin-2 in ECs results in angiogenesis defects**

(A) Schematic illustration of the Tamoxifen administration schedule for *Kindlin-2*$^{WT}$ and *Kindlin-2*$^{iΔEC(Cdh5)}$ mice. Tamoxifen was administered from P1 to P3, and retinas were analyzed at P6.

(B) Representative confocal images of flat-mounted retinas from P6 *Kindlin-2*$^{WT}$ and *Kindlin-2*$^{iΔEC(Cdh5)}$ pups stained with IsoB4 (red).

(C–D) Quantification of (B), showing reduced vascular coverage and branch points (n = 6/7 pups).

(E) Representative images of the angiogenic front in P6 *Kindlin-2*$^{WT}$ and *Kindlin-2*$^{iΔEC(Cdh5)}$ retinas stained with IsoB4. Higher-magnification images show the blunted morphology of tip cells.

(F) Quantification of (E), showing a reduced number of sprouts (n = 6/7 pups).

(G) Representative images of mouse aortic rings from *Kindlin-2*$^{WT}$ and *Kindlin-2*$^{iΔEC(Pdgfb)}$ mice cultured for 5 days in the presence of 5 μM 4-hydroxytamoxifen (4-OHT). Actin, phalloidin (green).

(H) Quantification of microvascular sprouting area relative to total ring area in (G) (n = 5/5 aortic rings).

(I) Western blot analysis of Kindlin-2 expression after transfection with siCtrl or si*KINDLIN-2* for 48 hrs.

(J) Quantification of Kindlin-2 knockdown efficiency in (I) (n=3 independent experiments).

(K) HUVECs transfected with siCtrl or si*KINDLIN-2* for 48 hrs were starved in 2% FBS for 8 hrs. A scratch wound was made in the monolayer, followed by treatment with 50 ng/ml VEGF or vehicle for 6 hrs. Cells were fixed and stained with phalloidin and visualized by confocal microscopy. White arrows indicate phalloidin$^+$ filopodia. Actin, phalloidin (green); nuclei, DAPI (white).

(L) Quantification of the number of phalloidin$^+$ filopodia from (K) based on 3 independent experiments.

(M) HUVECs transfected with siCtrl or si*KINDLIN-2* for 48 hrs were starved in 2% FBS for 8 hrs and treated with 50 ng/ml VEGF or vehicle, followed by 10 μM BrdU for 4 hrs. Cells were fixed, stained with DAPI, and visualized under a confocal microscope. BrdU (green); nuclei, DAPI (blue).

(N) Quantification of the percentage of BrdU$^+$ cells among total DAPI$^+$ cells in (M) (n=4 independent experiments).

(O) Expression of Integrin family proteins in human retinal endothelial cells (HRECs), HUVECs, human cortical microvessels endothelial Cells (HCMECs), and purified P5 mouse brain ECs. Data were analyzed from GSE 291008, GSE287078, GSE281250 and GSE94862.

Data are presented as mean ± SEM. * $P < 0.05$; ** $P < 0.01$; *** $P < 0.001$； ****$P < 0.0001$; two-tailed Student's t-test or one-way ANOVA followed by Tukey's multiple comparisons test.

Scale bars: 500 μm in (B), 200 μm in (G) and low magnification of (E), 100 μm in high magnification of (E), 50 μm in (M), 5 μm in (K).

**Figure S4. Kindlin-2 associates with Moesin in ECs**

(A) Co-IP analysis of Kindlin-2 and Moesin in HUVECs infected with Ad-Flag-Moesin$^{WT}$ or Ad-Flag-Vector. Kindlin-2 was pulled down with anti-Flag M2 beads and detected by western blot. Ad-Vector-infected cells served as control.

(B)Schematic illustration of the predicted binding model of WT Moesin and the N62A mutant.

(C) Expression of Moesin, Radixin, and Ezrin in HRECs, HUVECs, HCMECs, and purified P5 mouse brain ECs. Data were analyzed from GSE 291008, GSE287078, GSE281250 and GSE94862.

(D) Relative expression levels of Moesin in embryonic brain ECs and whole brain tissue at different developmental stages. Data were analyzed from GSE79306.

**Figure S5. Hypotonic treatment increases membrane tension and reduces VEGF signaling**

(A) Representative FRET ratiometric images of the MSS probe expressed in EA.hy926 cells treated with isotonic or hypotonic medium for a few seconds.

(B) Quantification of FRET efficiency from (A) (n = 30/29 cells from 3 independent experiments).

(C) HUVECs were starved in 2% FBS for 8 hrs, treated with isotonic, 200 mOsm, or 120 mOsm medium, then stimulated with Alexa 594-labelled VEGF (red) for 30 mins before fixation. VEGF accumulation was visualized by confocal microscopy. Actin, phalloidin (green); nuclei, DAPI (blue).

(D) Quantification of intracellular Alexa 594-labelled VEGF (red) spots from (C) (n=3 independent experiments).

(E) HUVECs starved for 8 hrs and treated with isotonic, 200 mOsm, or 120 mOsm medium, then stimulated with 50 ng/ml VEGF for 0, 5, 15, or 30 mins. Representative western blots show ERK phosphorylation in response to VEGF stimulation.

(F) Quantification of p-ERK levels at 5 mins after VEGF stimulation in (G) (n=3 independent experiments).

Data are presented as mean ± SEM. *$P < 0.05$; **$P < 0.01$; ***$P < 0.001$; two-tailed Student's t-test and one-way ANOVA followed by Tukey's multiple comparisons test.

Scale bars: 5 μm in (A), (C).

**Figure S6. Kindlin-2 deficiency alters membrane tension and VEGF signaling in a Moesin-dependent manner**

(A) Representative fluorescence lifetime images of HUVECs transfected with siCtrl, si*KINDLIN-2*, or si*KINDLIN-2* + si*MSN*. Flipper-TR probe (1.5 μM) was added to the medium and incubated at 37 °C for 15 mins before imaging by FLIM.

(B) Quantification of fluorescence lifetime from (A) (n = 27/23/27 cells from 3 independent experiments).

(C) HUVECs transfected as in (A) were starved for 8 hrs, stimulated with VEGF for 30 mins, fixed, and stained for VEGFR2 (green). Representative images show VEGFR2 vesicles; nuclei stained with DAPI (blue).

(D) Quantification of VEGFR2 vesicles per cell from (C) (n= 3 independent experiments).

(E) HUVECs transfected with siCtrl or si*MSN*-3'UTR followed by infection with Ad-Vector, Ad-Moesin$^{WT}$, or Ad-Moesin$^{N62A}$ were starved in 2% FBS for 8 hrs, stimulated with VEGF-A for 30 mins, fixed, and stained for VEGFR2 (green). Representative images show VEGFR2 vesicles; nuclei stained with DAPI (blue).

(F) Quantification of VEGFR2 vesicles per cell from (E) (n= 3 independent experiments).

Data are shown as mean ± SEM. *$P < 0.05$; **$P < 0.01$; ***$P < 0.001$; one-way ANOVA followed by Tukey's multiple comparisons test.

Scale bars: 5 μm in low magnification of (A), (C), and (E); 2.5 μm in high magnification of (A), (C), and (E).

**Figure S7. Kindlin-2–Moesin interaction in pathological angiogenesis**

(A) Uniform Manifold Approximation and Projection (UMAP) plots of scRNA-seq data (GSE216676) from P17 OIR mouse retina tissue showing various EC types present in the retina.

(B) Violin plot showing Kindlin-2 (encoded by *Fermt2*) expression levels across different EC types in the OIR mouse retina (GSE216676). Kindlin-2 is highly expressed in tip cells and tuft ECs.

(C) Representative retinal image showing colocalization of endothelial marker IsoB4 with ZsGreen expression mediated by AAV-BR1-ZsGreen 14 days after retro-orbital injection.

(D) Quantification of knockdown efficiency of AAV-sh*Msn* in isolated brain ECs.

Data are presented as mean ± SEM. ** $P < 0.01$, two-tailed Student's t-test.

Scale bars: 100 μm in (C).

**Table S1. Primers for qPCR**

| PRIMERS | SEQUENCE | |
|---|---|---|
| Mouse | Forward | Reverse |
| *Msn* | TCTTATGCCGTCCAGTCTAAGT | GGTCCTTGTTGAGTTTGTGCT |
| *Fermt2* | TGGACGGGATAAGGATGCCA | TGACATCGAGTTTTTCCACCAAC |
| *Atcb* | GGCTGTATTCCCCTCCATCG | CCAGTTGGTAACAATGCCATGT |

**Table S2. Antibodies**

| Target antigen | Vendor or Source | Catalog # | Working concentration |
|---|---|---|---|
| GAPDH | Utibody | UM4002 | 1:1000 (WB) |
| CD31 | Millipore | MAB1398Z | 1:100 (IF) |
| Flag | CST | 14793S | 1:1000 (WB) |
| goat anti-Rabbit | MULTI SCIENCES | GAR007 | 1:5000 (WB) |
| goat anti-Mouse | MULTI SCIENCES | GAM007 | 1:5000 (WB) |
| HA | CST | 2367S | 1:1000 (WB) |
| Kindlin-2 | Millipore | MAB2617 | 1:1000 (WB)<br>1:100 (IF)<br>1:50 (PLA) |
| Moesin | CST | 3150S | 1:1000 (WB)<br>1:100 (PLA) |
| VEGFR2 | CST | 2479S | 1:1000 (WB)<br>1:200 (IF) |
| p-ERM | CST | 3141S | 1:1000 (WB)<br>1:200 (IF) |
| Ki67 | Abcam | ab16667 | 1:100 (IF) |
| ERG | CST | ab92513 | 1:100 (IF) |
| Ter119 | R&D system | MAB1125 | 1:200 (IF) |
| Alexa Fluor 594-conjugated isolectinGS-IB4 | Invitrogen | I21413 | 1:200 (IF) |
| Alexa Fluor 488-conjugated isolectinGS-IB4 | Invitrogen | I21411 | 1:200 (IF) |
| p42/44 MAPK (Erk1/2) Antibody | CST | 9102S | 1:1000 (WB) |
| phospho-p42/44 MAPK (Erk1/2) Antibody | CST | 4370S | 1:1000 (WB) |
| Phalloidin | Invitrogen | A12349 | 1:300 (IF) |
| anti-BrdU | Abcam | ab6326 | 1:100 (IF) |

**Table S3. siRNA sequences**

| Oligonucleotides | SEQUENCE |
|---|---|
| Human si*KINDLIN-2*-1 | GCUUAAGCUGGUGGAGAAACUCGAUGUAA |
| Human si*KINDLIN-2*-2 | AACAGCGAGAAUCUUGGAGGCCCAUCAGA |
| Human si*MSN* | UCGCAAGCCUGAUACCAUUTT |
| Human si*MSN* 3'UTR | GCTAAATTGAAACCTGGAAT |
